# Plasticity and the role of mass-scaling in allocation, morphology and anatomical trait responses to above and belowground resource limitation in cultivated sunflower (*Helianthus annuus L.*)

**DOI:** 10.1101/504316

**Authors:** Yan Wang, Lisa A. Donovan, Andries A. Temme

## Abstract

- In the face of resource limitations, plants show plasticity in multiple trait categories, including biomass allocation, morphology and anatomy, yet inevitably also grow less. The extent to which passive mass-scaling plays a role in trait responses that contribute to increased potential for resource acquisition are poorly understood. Here we assessed the role of mass-scaling on the direction, magnitude and coordination of trait plasticity to light and/or nutrient limitation in cultivated sunflower (*Helianthus annuus*).
- We grew seedlings of ten sunflower genotypes for three weeks in a factorial of light (50% shade) and nutrient (10% supply) limitation in the greenhouse and measured a suite of allocational, morphological and anatomical traits for leaves, stems, fine roots, and tap roots.
- Under resource limitation, plants were smaller and more biomass was allocated to the organ capturing the most limiting resource, as expected. Traits varied in the magnitude of plasticity and the extent to which the observed response was passive (scaled with plant mass) and/or had an additional active component. None of the allocational responses were primarily passive. Plastic changes to specific leaf area and specific root length were primarily active, and adjusted towards more acquisitive trait values under light and nutrient limitation, respectively. For many traits, the observed response was a mixture of active and passive components, and for some traits the active adjustment was antagonistic to the direction of passive adjustment, e.g. stem height, and tap root and stem theoretical hydraulic conductance. Passive scaling with size played a major role in the coordinated response to light, but correcting for mass clarified that the active responses to both limitations were more similar in magnitude, although still resource and organ specific.
- Our results demonstrate that both passive plasticity and active plasticity can contribute to increased uptake capacity for limiting resources in a manner that is resource, organ and trait specific. Indeed, passive adjustments (scaling with mass) of traits due to resource stress extends well beyond just mass allocation traits. For a full understanding of plants response to environmental stress both passive and active plasticity needs to be taken into account.

## Introduction

The extent of plant trait adjustment in response to a changed environment is generally considered as the plant’s phenotypic plasticity (Valladares, Gianoli, & Gomez, 2007; Nicotra et al., 2010). According to theory, this plasticity serves to optimize/maximize the uptake of the most limiting resource (Bloom, Chapin, & Mooney, 1985; Gedroc, McConnaughay, & Coleman, 1996; Shipley & Meziane, 2002; Poorter et al., 2012). For example, increased mass allocation to leaves under shade, and to roots under nutrient limitation, alleviates some of the stress caused by the resource limitation (Shipley & Meziane, 2002; Sugiura & Tateno, 2011). However, plants are inevitably smaller under resource stress, raising the question of the role of mass-scaling in individual and coordinated traits shifts in response to resource limitation (McConnaughay & Coleman 1999; Shipley & Meziane, 2002; Weiner 2004; Poorter et al. 2012; Osnas, Lichstein, Reich, & Pacala, 2013; Reich, 2014 and 2018).

There is a long tradition of investigating mass-based scaling of biomass allocation to different plant parts (Weiner 2004). Generally, smaller plants allocate proportionally more mass to leaves than to roots (Poorter et al., 2012). Trait responses that scale with mass, which have been variously called allometric scaling, “passive” plasticity, or “apparent” plasticity, could predispose the plant to align its traits with resource availability and demand (McConnaughay & Coleman, 1999; Weiner 2004; van Kleunen & Fisher, 2005; Nicotra et al., 2010; Poorter et al., 2012). However, there is evidence that additional “active” or “true” plasticity in biomass allocation can further increase the capacity for acquiring the most limiting resource (McConnaughay & Coleman, 1999; Shipley & Meziane, 2002; Poorter et al. 2012). There is also evidence of active plasticity in other traits related to resource uptake. For example, under low light conditions greater plant height and greater specific leaf area for a given plant mass aids in light uptake (Rice & Bazzaz, 1989; Freschet, Swart, & Cornelissen, 2015; Freschet, Violle, Bourget, Scherer-Lorenzen, & Fort, 2018; Reich, 2018). Thus, both passive and active trait adjustments combine in the realized response to resource stress.

The role of passive and active contributions to observed responses can be investigated by including analyses that correct responses for plant size (Poorter et al., 2012; Reich, 2018). In this framework, passive adjustments in traits associated with size cannot be a priori regarded as or ruled out as adaptive (Nicotra et al., 2010; Poorter et al., 2012 and 2019). If traits responses that are consistent with greater ability to take up the most limiting resource have both passive and active components, it will be important to consider both the magnitude and alignment of both components when evaluating evidence for functional and putatively adaptive responses.

Among plant traits, anatomical traits are often overlooked due to time and budget constraints. However, variation in anatomical traits underlies or contributes to variation in morphological and physiological traits that have received more attention (Kong et al., 2014; Scoffoni et al., 2015; John et al., 2017). For example, palisade parenchyma thickness is positively correlated to leaf thickness (Catoni, Gratani, Sartori, Varone, & Granata, 2015) and photosynthetic rate (Chatelet, Clement, Sack, Donoghue, & Edwards, 2013). Additionally, a thicker cortex could provide a relative larger site for mycorrhizal infection and higher resource uptake in thicker roots, especially for the arbuscular mycorrhiza (Kong et al., 2014). Root cortex thickness, due to the size of cortical cells (Eissenstat & Achor, 1999) strongly affects fine root diameter (Gu, Xu, Dong, Wang, & Wang, 2014; Guo et al., 2008), and a wider stele and/or xylem conduit greatly affects hydraulic conductivity (McElrone, Pockman, Martínez-Vilalta, & Jackson, 2004; Rico, Pitterman, Polley, Aspinwell, & Fay, 2013; Tyree & Ewers, 1991). Thus, xploring how passive and active responses in anatomical traits align with those of other traits will enhance our understanding of how plants adjust to changing environmental conditions from tissue, to organ, to architecture.

Across species, plant functional traits are thought to form a spectrum of resource use strategies from fast to slow (Wright et al., 2004; Fortunel, Fine, & Baraloto, 2012; Reich, 2014; Díaz et al., 2016). The “whole plant economic spectrum” suggests that values of traits related to resource use, including mass allocation, morphology and transport, should be coordinated across all organs and resources, with more resource conservative traits associated with adaptation to light and nutrient limited habitats (Reich, 2014). For closely related species, there is mixed evidence of correlated evolution among some traits related to resource acquisition and processing (Mason & Donovan, 2015; Bowsher, Mason, Goolsby, & Donovan 2016; Pilote & Donovan, 2016; Medeiros, Burns, Nicolson, Rogers, & Valvarde-Barrantes, 2017; Muir, Conesa, Roldán, Molins, & Galmés, 2017). However, much remains unclear about how the coordinated traits across ‘whole plant economic spectrum’ relate to trait plasticity in response to resource limitations (Reich, 2014; Anderegg et al., 2018; Agrawal, 2020). Thus, it is interesting to assess the coordination of responses across organs and a broad range of traits in terms of the role of passive and active plasticity.

To add to our understanding of plant responses to resource limitation, we examined trait responses to light and nutrient limitation of traits across different trait categories (biomass allocation, morphology and anatomy) and organs (leaf, stem and root) in cultivated sunflower. Prior research has shown strong plastic responses to resource limitation and other environmental factors in *H. annuus* (Rico, Pitterman, Polley, Aspinwell, & Fay, 2013; Donovan, Mason, Bowsher, Goolsby, & Ishibashi, 2014; Bowsher et al., 2017; Masalia, Temme, Torralba, & Burke, 2018; Temme et al., 2019). Specifically, we sought to answer the following questions:

1. How do mass allocation, organ morphology, and anatomy change with above and below ground resource limitation, and what role does size scaling of traits play in this?
2. How do traits compare for magnitude of plasticity and what role does size scaling of traits play in this?
3. Do traits show a coordinated shift due to resource limitation across all organs and what role does size scaling of traits play in this?

## Material and Methods

### Experimental design

To address these questions, we selected a set of 10 cultivated sunflower genotypes (Supplemental Table S1), varying broadly in biomass based on prior work, from a larger diversity panel used for genomic dissection of traits (Mandel, Dechaine, Marek, & Burke, 2011; Nambeesan et al., 2015; Masalia, Temme, Torralba, & Burke, 2018). We conducted a factorial design of two nutrient treatments (rich and poor) and two light treatments (sun and shade) at the Botany greenhouses of The University of Georgia, Athens GA, USA, in March 2018. Achenes were sown in seedling trays and allowed to grow for seven days, after which each seedling was transplanted to 5 liter (1.3 gallon) pot filled with a 3:1 sand:calcinated clay mixture (Turface MVP, Turface Athletics, Buffalo Grove, IL). Pots were arranged in a split plot design of 6 replicate blocks. The light treatment was applied as the whole-plot factor, with 2 sub-plots in each plot randomly assigned to unshaded or 50% shade generated with high density woven polyethylene cloth (Supplemental Figure S1). Within each subplot, two pots of each genotype were randomly distributed and supplied with either 40g or 4g fertilizer (Osmocote Plus 15-9-12 with micronutrients, Scotts, Marysville, OH, USA), totaling 240 pots (plants). Greenhouse temperature controls were set to maintain 18–24 °C, and natural sunlight was supplemented with sodium halide lighting (20-25 μmol m^−2^ s^−1^) to maintain a 15/9-h photoperiod.

### Plant harvest and trait measurements

Plants were harvested 3 weeks after transplanting (4 weeks after germination). At harvest, stem height (ST-Hgt, from soil surface to top of apical meristem) and stem diameter (ST-Dia, midway between cotyledons and first leaf pair) were measured. Plants were separated into root, leaf (including cotyledons), and stem (including bud if present—rarely) for biomass and other measurements. We assigned all measured traits to one of three categories (allocational, morphological, anatomical) in order to compare the relative magnitude of adjustments to these broad categories. While we believe our assignment of traits to categories is defensible, we acknowledge that this is somewhat arbitrary and that different groupings could influence results associated with comparisons among categories.

Each replicate plant was sampled for leaf, stem and root tissue for anatomical traits. One recently matured fully expanded leaf was sampled for a 1×0.5 cm rectangle cut out of the leaf center. The stem was sampled for a 5 mm length segment centered between the cotyledon and the first leaf pair. The root was sampled for both tap root and fine root tissue. For the tap root, a 1 cm segment was cut 4 cm below the root/stem junction. For a single lateral root attached to the tap root near to the root/stem junction with an intact root tip, a 1 cm fine root segment was cut 2 cm from the apex of the root. All tissue subsamples were weighed for fresh mass and then fixed in formalin–acetic acid-alcohol solution, FAA (50% ethanol (95%), 5% glacial acetic acid, 10% formaldehyde (37%) and 35% distilled water).

Fixed subsamples were processed for anatomy at the University of Georgia Veterinary Histology Laboratory. Each sample was embedded and gradually infiltrated with paraffin, sliced with a sledge microtome, mounted to a slide, and stained with safranin and fast green dye. Slides were imaged with a camera mounted Zeiss light microscope using ZEN software (Carl Zeiss Microscopy, Oberkochen, Germany). For leaf, stem, and fine and tap roots, the dimensions of anatomical features were traced using Motic Images Advanced 3.2 software (Motic Corporation, Xiamen, China). The organs dimensions included leaf thickness LF-Th), fine root diameter (FR-Dia) and tap root diameter (TR-Dia). The tissue dimensions included leaf palisade thickness (LF-PT), Leaf spongy thickness (LF-ST), stem cortex thickness (ST-CT), Stem xylem thickness (ST-XT), tap root cortex thickness (TR-CT), tap root stele diameter (TR-SDia), fine root cortex thickness (FR-CT), and fine root stele diameter (FR-SDia). The theoretical hydraulic conductivity (Ks, kg·;s^−1^·m^−1^·MPa^−1^) per vascular area for leaf (LF-Ks), stem (ST-Ks), fine root (FR-Ks), and tap root (TR-Ks) tissue was calculated (Sperry, Hacke, & Pittermann, 2006), based on the Hagen-Poseuille equation (Tyree & Ewers, 1991): 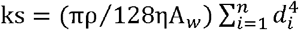. where ρ is the density of water (988.3 kg·m^−3^ at 20 °C); η is the viscosity of water (1.002×10^−9^ MPa·s at 20 °C); A_w_ is the stele (vascular) cross-section area, d is the diameter of the i^th^ vessel and n is the number of conduits in the xylem.

After anatomical samples were collected, the remaining plant material was used to determine morphological traits. The leaf, stem, and fine and tap roots for each plant were weighed for fresh mass. Root tissue, stem tissue, and the most recently fully expanded leaf was scanned at 300 dpi with an Epson Expression1680 scanner (Seiko Epson Corporation, Nagano-ken, Japan) and saved as a TIF image. Total root length and volume of scanned fine root, tap root and stem, as well as leaf area were measured using WinRhizo (v. 2002c, Regent Instruments, Quebec, Canada). Subsequently, the leaf, stem, and fine roots and tap roots were dried at 60°C for 48 h and weighed. Specific leaf area (LF-SLA, cm^2^·g^−1^) was calculated as the ratio of leaf area to leaf dry mass. Specific root length (m·g^−1^) was calculated as the ratio of root length to root dry mass for the tap root and fine roots (TR-SRL, FR-SRL, respectively). Tissue density (g·cm^−3^) was calculated as the ratio of dry mass to volume for stem, tap toot, and fine roots (ST-Den, TR-Den, FR-Den, respectively). Leaf dry matter content, measured as leaf dry mass divided by leaf fresh mass, was used as a proxy for leaf tissue density (LF-Den) (Kramer-Walter et al., 2016; Wilson, Thompson, & Hodgson, 1999).

For allocational traits, total plant dry mass was calculated as the sum of all plant parts, including the subsamples for anatomical and morphological traits. The fresh biomass of subsamples for anatomical analysis was converted to dry biomass based on the ratio of fresh/dry biomass of for the morphological traits. The mass fractions for each tissue were calculated as proportions of total plant dry mass (g·g^−1^).

### Data analysis

The statistical analysis for the phenotypic data was performed using R v3.5.1 (R Core Team). To obtain genotype means from our split plot design, a mixed effects model was fitted using the package *lme4* (Bates et al., 2018) with genotype, light and nutrient level and all their interactions as fixed effects and light treatment within block as random factor. Least-square (LS) means of all trait values without random factor were estimated from this model using the R package emmeans (Lenth et al., 2018). To test the effect of genotype and treatment on measured traits we fitted a less expansive mixed effects model with genotype, light and nutrient level as well as the interaction between light and nutrient level as fixed effects (following Freschet, Violle, Bourget, Scherer-Lorenzen, & Fort, 2018) and light treatment within block as random factor. From this model, fixed effects were then tested using a Walds Chi-square test in a type III ANONA using the package car (Fox et al., 2018). As we were interested in the interactive effects among genotype, light and nutrient supply on the proportional changes in functional traits, rather than on their absolute changes, we performed all analyses on log-transformed data (Freschet, Swart, & Cornelissen, 2015). For each trait, differences among treatments were tested using Tukey’s HSD (p=0.05) corrected for multiple comparisons. We then estimated the influence of plant size on the significance of nutrient and light limitation effects on traits by adding (log-transformed) plant biomass as a fixed factor to both models and recalculating means and significance (Ryser & Eek, 2000; Wahl, Ryser, & Edwards, 2001).

To quantify the plastic response of each trait to each resource limitation treatment, we calculated the relative distance plasticity index (RDPI, Valladares, Sanchez-Gomez, & Zavala, 2006, Scoffoni et al., 2015) as 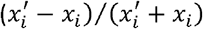, where *x*_*i*_ and 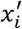 are the mean trait values of genotypes grown under control (high-light and high-nutrient) and resource limited condition, respectively. Additionally, the RDPI for each trait in each resource limitation treatment was recalculated after correcting for plant size (total biomass). Significant values of RDPI (difference from zero, no plasticity) were determined using T-test on genotype averages.

To determine major axes of variation across multiple traits and identify whether there were concerted trait adjustments to limitation in above or belowground resources, we conducted a principal component analysis (PCA) on the trait data before and after correcting for size. Differences between treatments were tested using, Bonferroni corrected, Hotellings-t test on the first two principal components. Data visualizations were made using *ggplot2* (Wickham et al., 2018).

## Results

### How do plant mass allocation, organ morphology, and anatomy respond to above and below ground resource limitation, and what role does size scaling of traits play in this?

Across all genotypes, total plant biomass decreased substantially in response to light and nutrient limitation (Figure 1 and Table 1). The biomass response to light limitation (54.7% decline) was stronger than the response to the nutrient limitation (24.7% decline), and there was no interaction of light and nutrient limitation on biomass when combined (67.1%% decline). Of the 31 allocational, morphological and anatomical traits measured, all but two responded to at least one resource limitation (Supplemental Table S2). Responses to light limitation were more prevalent than responses to nutrient limitation, and there were relatively few interactive effects of combined light and nutrient limitation.

**Figure 1.**
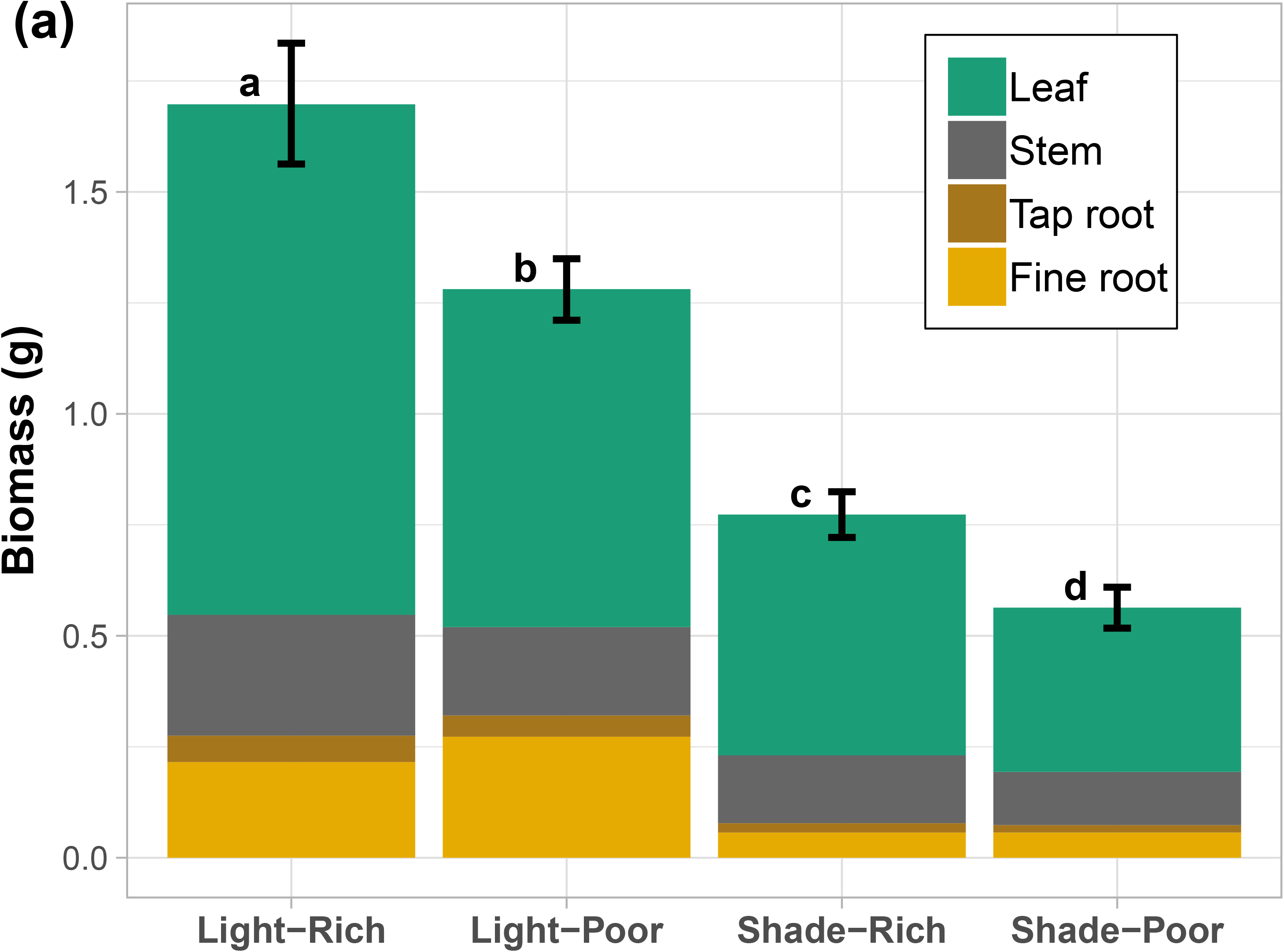
Biomass and mass allocation of leaf, stem, tap root, and fine root tissue across a factorial of light and nutrient limitation. Light/Shade, direct sun vs 50% shade. Rich/poor, high nutrients vs 10% nutrient concentration. Whole plant biomass (mean±SE) of 10 genotypes (estimated marginal mean from the ANOVA, based on 5-6 replicates per genotype), stacked by average tissue contribution. Different lower-case letters represent significant (p<0.05) Tukey post hoc differences between treatments for whole plant biomass.

For allocational traits, resource limitations affected all 6 traits, but traits were affected in contrasting ways by light and nutrient limitation. The RatioLF-FRmass (ratio of leaf mass to fine root mass) increased under light limitation due to increased LF-MF (leaf mass fraction) and decreased FR-MF (fine root mass fraction) (Figures 2b-d and Table 1). Conversely, RatioLF-FRmass decreased under nutrient limitation, due to decreased LF-MF and increased FR-MF (Figure 2f). A significant interaction of light and nutrient limitation was found for LF-MF, ST-MF (stem mass fraction), FR-MF, and RatioLF-FRmass. The RatioSLA-SRL (ratio of LF-SLA (specific leaf area) to FR-SRL (fine root specific root length)), which additionally considers surface area per unit mass of acquisitive organ, paralleled that of the RatioLF-FRmass, with a trend of increasing under light limitation and decreasing under nutrient limitation (Figure 2e).

**Figure 2.**
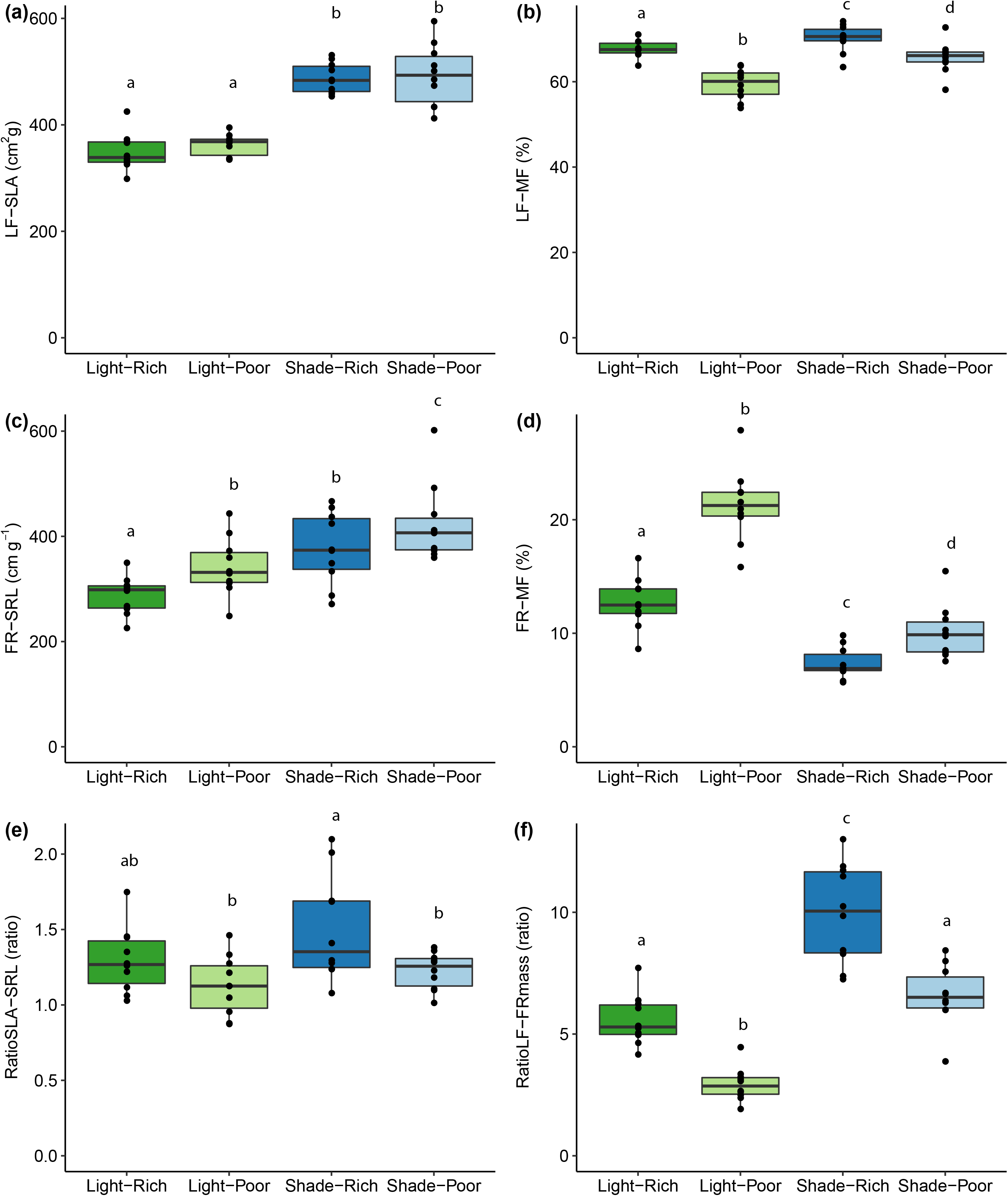
Leaf and fine root morphology across a factorial of light and nutrient limitation. Light/Shade, direct sun vs 50% shade. Rich/poor, high nutrients vs 10% nutrient concentration. **(a)** specific leaf area (LF-SLA, cm^2^ g^−1^), **(b)** leaf mass fraction (LF-MF, %), **(c)** specific root length (FR-SRL, cm g^−1^), **(d)** fine root mass fraction (FR-MF, %), **(e)** ratio of leaf area to root length (RatioSLA-SRL), **(f)** ratio of leaf mass to fine root mass (RatioLF-FRmass). Points indicate genotype (n=10) mean (n=5-6) at a given treatment. Boxplots show distribution of values. Different letters represent significant (p<0.05) Tukey post hoc differences between treatments for each trait.

For morphological traits, light limitation affected 11 of the 12 traits, all except for FR-Dia (fine root diameter) (Table 1). The responses led to increases in acquisitive values for resource acquiring traits such as LF-SLA (Figure 2a), FR-SRL (Figure 2c), TR-SRL (tap root specific root length), and ST-Hgt (stem height). There were also strong decreases in organ dimensions, e.g., LF-Th (leaf thickness), ST-Dia (stem diameter), and TR-Dia (tap root diameter). LF-Den (leaf density), ST-Den (stem density), and FR-Den (fine root density) also declined. Contrasting with the effect of light limitation, nutrient limitation affected only four out of 12 morphological traits (Table 1), leading to decreased ST-Dia and FR-Dia, and increased FR-SRL and TR-Den (tap root density).

For anatomical traits, limiting resources affected 12 of the 13 traits, and responses again differed by limiting resource (Figure 3 and Table 1). Light limitation affected anatomical traits of all three organs. Aboveground, light limitation decreased LF-PTh (leaf palisade parenchyma layer thickness), ST-CTh (stem cortex thickness), ST-VTh (stem vascular bundle thickness), and ST-XTh (stem xylem thickness). Belowground, light limitation decreased TR-CTh (tap root cortex thickness) and TR-SDia (tap root stele diameter). Vascular tissue adjustment to light limitation led to changes in theoretical hydraulic conductivity, with decreased LF-Ks (leaf hydraulic conductivity) but increased ST-Ks (stem hydraulic conductivity). In contrast to the effects of light limitation, nutrient limitation predominately affected fine root anatomy, leading to decreased FR-CTh (fine root cortex thickness) and FR-SDia (fine root stele diameter).

**Figure 3.**
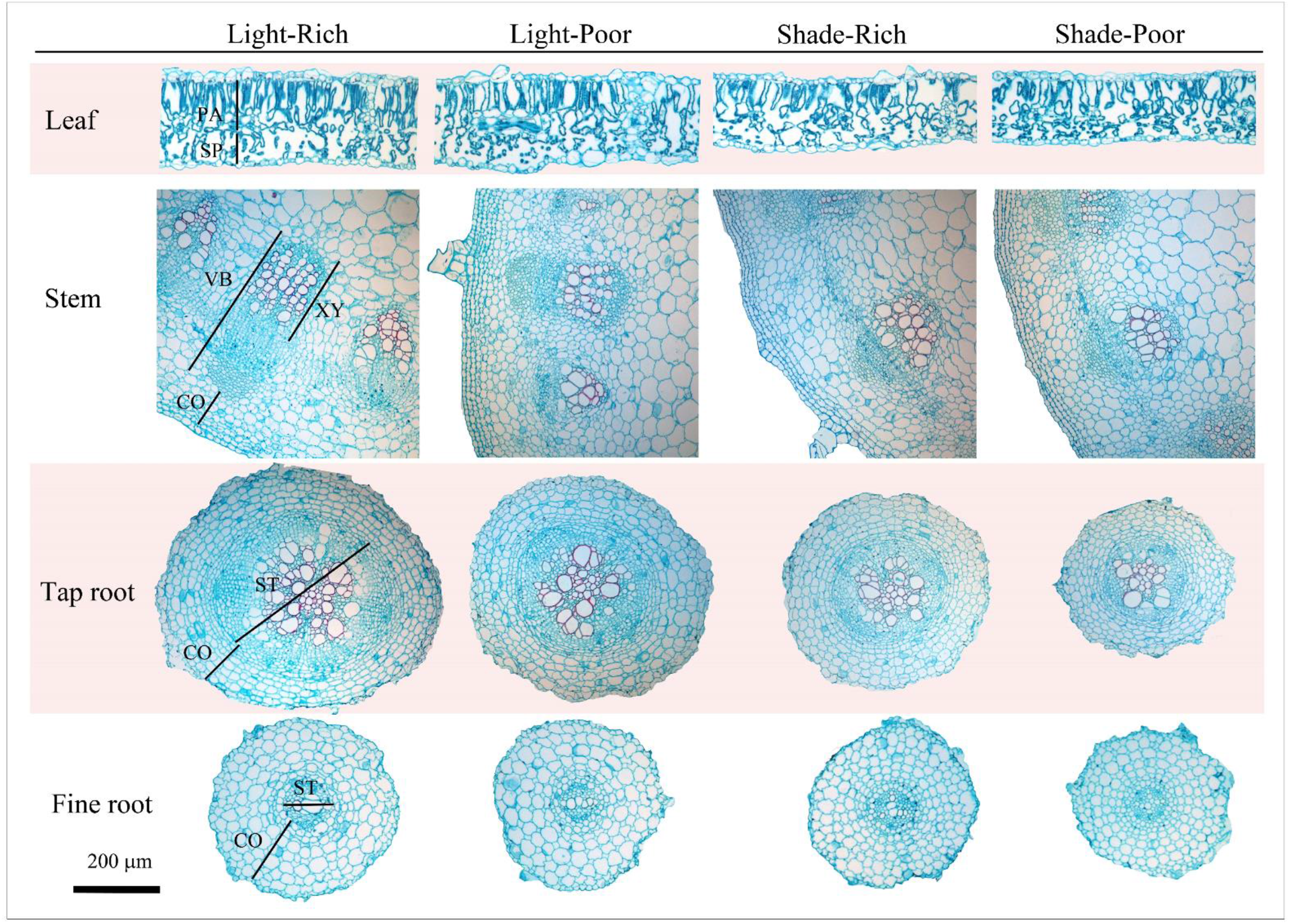
Typical anatomical structure of leaf, stem, tap root and fine root of genotype HA‐across a factorial of light and nutrient limitation. Light/Shade, direct sun vs 50% shade. Rich/poor, high nutrients vs 10% nutrient concentration. PA, palisade tissue; SP, spongy tissue; CO, cortex; VB, vascular bundle; XY, xylem; ST, stele.

To assess the role of passive mass-based scaling in the responses of resource limitations, trait responses were reanalyzed with mass as a covariate in the model (Table 1, Supplemental Figure S3), effectively comparing treatments again for trait values of plant of the same size. Contrasting the results of the two models (including mass as covariate in Table 1), we designated the observed response as being primarily passive (scaling with mass), primarily active (adjusted independent of mass scaling), or a mixture of both (Table 1).

For six traits, the responses to resource limitation were primarily passive responses, with effects no longer evident when mass was included as covariate). This included two morphological traits (FR-Den and TR-SRL) and four anatomical traits (LF-Ks, ST-CTh, ST-XTh, and TR-CTh). For these traits, the observed response to light limitation primarily was associated with smaller plant size.

For 13 traits, the responses were primarily active, with effects not changing when mass was included in the model, indicating that the response was still evident after correcting for plant size. This included 3 allocational traits (ST-MF, FR-MF, and TR-MF), six morphological traits (LF-Th, LF-SLA, LF-Den, ST-Den, FR-Den, and FR-SRL) and four anatomical traits (LF-PT, LF-ST, FR-CT, and FR-SD). For these traits, resource limitation effects were not associated with smaller plant size.

For 11 traits, the responses were a mixture of passive and active responses to resource limitation, with the effects differed somewhat when mass was included in the model, indicating that scaling with mass accounted for a portion of the response. This included 3 allocational traits (LF-MF, RatioLF-FRmass, and RatioSLA-SRL), four morphological traits (ST-Dia, ST-Hgt, TR-Dia, and TR-Den) and four anatomical traits (ST-VTh, ST-Ks, TR-SDia, and TR-Ks).

### How do traits compare for magnitude of relative plasticity and what role does size scaling of traits play in this?

The magnitude and direction of trait plasticity in response to resource limitation was assessed with a relative distance plasticity index (RDPI, Figure 4a). The RDPI for at least one of the treatments was significantly different from control for all of the traits expect FR-Ks. Visually comparing RDPI per trait category showed that allocational traits had the greatest plastic responses, followed by morphology, and then anatomy. Although the treatment that combined light and nutrient limitation (“shade-poor” in Figure 4a) resulted in the largest decline in biomass, it was not consistently the treatment that induced the greatest magnitude of plasticity across all of the traits.

**Figure 4.**
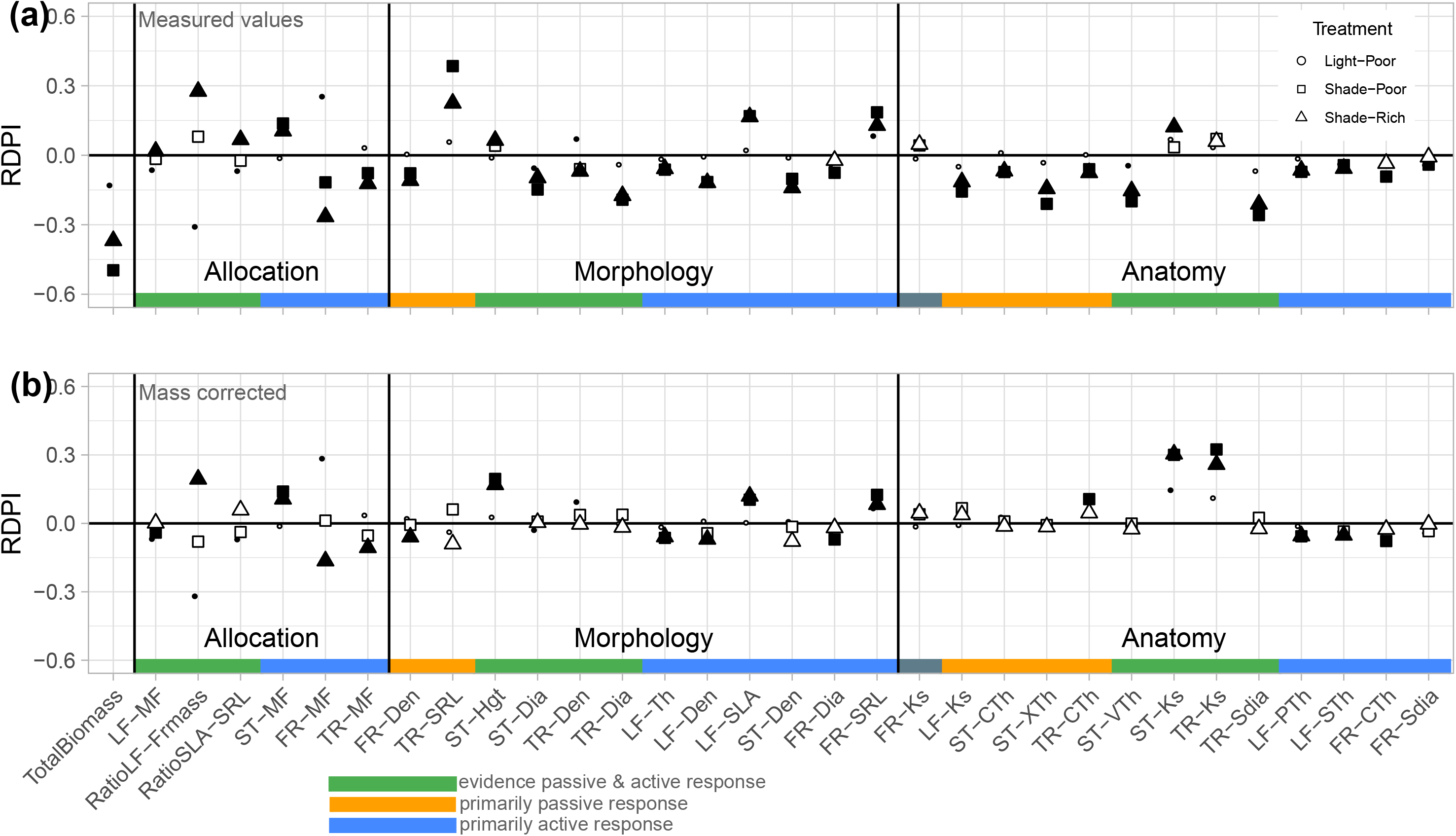
Trait plasticity in response to resource limitation. Points indicate the average (n=10) relative distance plasticity index (RDPI) in response to light limitation (triangles, Shade-Rich) or nutrient limitation (circles, Light-Poor), or combined (squares, Shade-Poor). Symbol fill represents a T-test significance (p<0.05) of RDPI being different from zero (filled symbols) or not (open symbols). Traits are ordered based on their treatment response, primarily passive (scaling with mass), primarily active (adjusted independent of mass), or a combination thereof. **(a)** RDPI values taken from base measurements. **(b)** RDPI values when correcting trait values for biomass. Trait abbreviations as in Table 1.

When trait responses to resource limitation were corrected for plant mass (Figure 4b), the extent to which RDPI values were affected was variable. Since this RDPI analysis is based on genotype means instead of individual plants there are slight differences in the extent of active vs passive responses though results broadly line up with the prior analysis (Table 1). For traits where responses were designated as primarily passive (Table 1), correcting for plant mass resulted in substantially lower RDPI (e.g., comparing RDPI in 4b to 4a for TR-SRL, LF-Ks, and ST-XT). For traits where responses were designated as primarily active, correcting for plant mass had little effect on RDPI (e.g., comparing 4b to 4a for ST-MF, FR-MF, LF-SLA, and LF-PT). For traits where responses were designated as a combination of active and passive, correcting for plant mass had a variable effect on RDPI. For some traits, RDPI was reduced for at least one treatment when mass was factored out (e.g., LF-MF, RatioLF-FRmass, and ST-Dia) indicating that the passive and active components that response were complementary in direction, resulting in a greater overall magnitude. However, for a few traits such as ST-Hgt, ST-Ks and TR-Ks, correcting for plant mass resulted in a greater RDPI, indicating active and passive responses to be in opposite directions, or antagonistic (Supplemental Figure S2).

### Do traits show a coordinated shift due to resource limitation across all organs and what role does size scaling of traits play in this?

When the genotypes means for the observed traits in all three traits categories (allocation, morphology, and anatomy) and all treatments were included in a PCA the correlations among many of the traits become evident. PC1 and PC2 explained 34.4% and 13.4% of the variation, respectively (Figure 5a). Light limitation resulted in a shift of a key set of traits to be more acquisitive along the first axis, with higher LF-SLA, TR-SRL, and FR-SRL under shade associated with lower values of other morphological traits (LF-Den, ST-Dia, TR-Dia, TR-SDia) and lower anatomical trait values (ST-XTh, ST-VTh). Nutrient limitation resulted in a smaller shift in traits, oriented more on PC2, and dominated by higher TR-MF, TR-Den, and TR-Ks and lower RatioSLA-SRL, ST-Hgt, FR-Dia, FR-SDia (Suplemental Figure S4).

**Figure 5.**
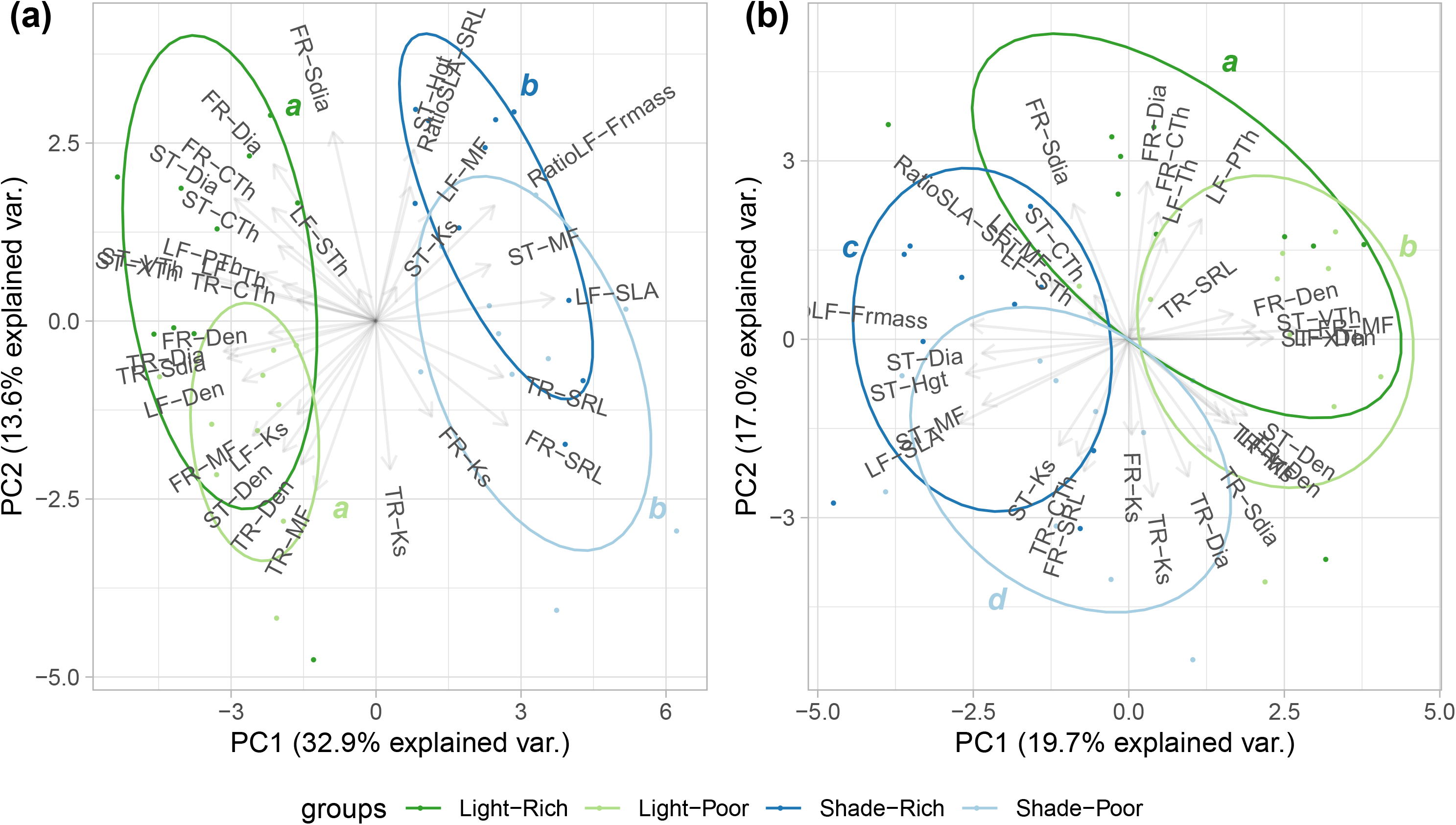
Principal component analysis (PCA) of leaf, stem and roots traits in response to resource limitations. Light/Shade, direct sun vs 50% shade. Rich/poor, high nutrients vs 10% nutrient concentration. Panels represent **(a)** trait values as measured, **(b)** trait values after correcting for size scaling (allometric effects). Different letters indicate significant Hotellings-T test, after Bonferroni correction, between treatments on the first two principal components. Trait abbreviations and units as in Table 1.

When the trait values corrected for mass were included in a PCA, reflecting primarily active responses, the correlations among traits changed and less of the variation was explained (PC1 and PC2 explained 19.7% and 17.0% of the variation, respectively). Responses to light and nutrient limitation were more similar in scale, although still resource specific (Figure 5b). Thus, while the passive mass-effect of light limitation might be greater, the active responses to above and below ground resource limitation were similar in magnitude.

## Discussion

Here we assessed the direction, magnitude, and coordination of cultivated H. annuus responses to light or/and nutrient limitation for traits in multiple categories (allocation, morphology and anatomy) and across all organ (leaves, stem, and roots), using plants harvest at a similar age and developmental stage. We found that more biomass was allocated to the organs acquiring the most limiting resource (roots for nutrient stress and leaves for shade) and that there were additional morphological and anatomical trait adjustments that generally led to more acquisitive trait values (e.g. higher LF-SLA in shade and higher FR-SRL in nutrient stress). We explored the role of mass-scaling in these responses by looking at the effect of including whole plant dry mass as a covariate to correct for plant size. The trait responses to resource limitations varied in the extent to which they were primarily passive (i.e., associated with lower biomass of stressed plants), primarily active (not or only marginally associated with lower biomass of stressed plants) or a combination thereof. The variable magnitude and direction of active plasticity played an appreciable role in shaping the individual trait responses and the overall coordinated response that was unique to each resource limitation.

### Allocation, morphology, and anatomy responses to above and below ground resource limitations

Light limitation resulted in increased relative investment in aboveground plant parts and nutrient limitation resulted in increased relative investment in belowground parts, consistent with the expectation that plasticity serves to optimize/maximize the uptake of the most limiting resource (Figure 1 and Table 1) (Bloom, Chapin, & Money, 1985; Shipley & Meziane 2002; Poorter et al., 2012). This allocation pattern has been confirmed in other species and growth forms (e.g., grasses, Siebenkäs, Schumacher, & Roscher, 2015; shrubs, Valladares, Wright, Lasso, Kitajima, & Pearcy, 2000; trees, Reich, Tjoelker, Walters, Vanderklein, & Bushena, 1998, Poorter et al. 2012; Kramer-Walter & Laughlin, 2017). Additionally, morphological and anatomical adjustments resulted in a higher LF-SLA and FR-SRL, under light and nutrient limitations, respectively, decreasing the ratio of mass invested to resource uptake potential (Figures 2,3). Anatomical adjustments in leaf and root, such as decreased LF-PTh and FR-CTh, also reduced tissue metabolic and maintenance costs (Guo, Cao, & Xu, 2006, Galindo-Castañeda, Brown, & Lynch, 2018; Jaramillo, Nord, Chimungu, Brown, & Lynch, 2013). Thus, for *H. annuus*, greater capacity for resource acquisition was coupled with reduced costs, and this was achieved by a combination of passive and active plasticity.

The traits varied in the extent to which passive plasticity played a role in explaining the observed response to resource limitations (Figure 4 and Table 2). The response to both light and nutrient limitation appeared primarily passive for several morphological and anatomical traits, but not for any of the allocational responses. And, the extent of passive adjustment differed by resource. Similar to Poorter et al., 2012, we found that responses to light limitation were more often passive than for nutrient limitation. For many traits there was evidence of a mixture of passive and active responses, which could be complementary or antagonistic and thus influence the magnitude of the overall/observed response (Figure 4, Supplemental figure S2). However, after correcting for plant mass, treatment effects remained significant for all of the allocation traits and many morphological and anatomical traits, such as LF-SLA (Freschet, Violle, Bourget, Scherer-Lorenzen, & Fort, 2018; Reich, 2018; this study), FR-SRL, LF-PTh and FR-CTh (this study), indicating active adjustments of these key traits for obtaining the most limited above or belowground resource.

### Magnitude and direction of plasticity

Consistent with other resource limitation studies, the observed plasticity of mass allocational traits was largest, followed by morphological traits (Kramer-Walter & Laughlin, 2017, Valladares, Wright, Lasso, Kitajima, & Pearcy, 2000), and smallest in anatomical traits (Catoni, Gratani, Sartori, Varone, & Granata, 2015; Cai, Ji, Yan, Jiang, & Fang, 2017; Xu, Wang, Liu, Lu, & Guo, 2015). It should be noted that differences in the extent of plasticity for different categories of traits may be species specific or based on the traits included. For example, oak (*Quercus robur*) seedlings were more plastic in physiological traits under shade, yet beech (*Fagus sylvatica*) was more plastic in morphological traits (Valladares et al., 2002). Differences among trait categories for the magnitude of plasticity may reflect an inherent hierarchy originating from the internal component traits of any specific organ. For example, leaf thickness is highly correlated with palisade parenchyma (Scoffoni et al., 2015) and root diameter with cortex and/or stele thickness (Kong et al., 2014). Thus, small shifts in individual component anatomical traits could add up to larger shifts in morphological traits which in turn affect allocational traits.

For traits where there was evidence of a mixture of passive and active plasticity, the alignment of both components (i.e. complementary or antagonistic in direction) affected the direction and magnitude of the observed response (Figure 4, Supplemental Figure S2). For RatioLF-FRmass, the passive and active responses to light limitation combined to enhance the relative investment in leaves, increasing the capacity for light acquisition. For other traits, correcting for mass revealed substantive active plasticity that was antagonistic to passive plasticity. For example, the active plasticity for ST-Hgt, and ST-Ks and TR-Ks was of greater magnitude than the observed response. The active adjustments counteracted the passive scaling with mass and resulted in shaded plants with higher conducting capacity (TR-Ks, ST-Ks) as well as high axial transportation distance and total transpiring surface (ST-Hgt and LF-SLA). This suggests that a coordinated increase in axial root and stem hydraulic transport offset greater resistance due to a longer transportation distance (Plavcová & Hacke, 2012), facilitating efficient movement of carbon to roots and nutrients and water to leaves (Maurel, Simonneau, & Sutka, 2010; Rodríguez-Gamir, Primo-Millo, & Forner-Giner, 2016; Wahl, Ryser, & Edwads, 2001). Thus, both passive and active trait adjustments to resource limitation serve to increase resource uptake capacity and maintain (optimal) functioning of altered organs.

Where passive and active trait adjustments are antagonistic this could indicate a departure from “safe” trait values under non-limiting conditions. For example, increased ST-Ks may come at the cost of increased risk of xylem embolism in shaded plants (Tyree & Zimmermann, 2002). The anatomical dataset collected in this work provides an excellent resource for anatomical water flow models (Couvreur et al., 2018) to further shed light on the consequences of these anatomical trait adjustments for plant hydraulics and how plants balance resource uptake demands with stress safety margins.

### Coordinated trait shifts

Resource limitation led to a coordinated shift amongst many correlated traits, with an overall shift towards greater potential for resource acquisition, consistent with other resource manipulation studies for commonly measured traits (Freschet, Swart, & Cornelissen, 2015). The coordinated trait shifts differed by limiting resource, with light limitation dominated by higher LF-SLA and associated with lower FL-Den and shifts in other stem and tap root traits, while nutrient limitation dominated by thinner FR-SDia, FR-Dia, and TR-MF and associated with other root and stem traits. (Figure 5 and Table 1). Correcting traits for plant size altered the magnitude of responses and relationships among traits, and revealed that the active effects of light and nutrient limitation were still resource specific, but more similar in magnitude.

The coordinated trait plasticity in response to resource limitations was generally a resource specific shift in traits towards more acquisitive strategy, that was not evident across all organs. It is interesting to note that these patterns are not consistent with the “whole plant economic spectrum” expectation of correlated evolution of conservative traits across all organs and resource limitations (Reich 2014, Díaz et al., 2016). This plasticity is an example of extensive responses to resource limitations and other abiotic or biotic factors that undoubtedly contribute to variation captured in broader surveys of plants specialized in different habitats (Anderegg et al., 2018). As such, these findings contribute to ongoing efforts to understand the scale dependence of trait covariation and the role that trait plasticity plays at both ecological and evolutionary scales.

## Supporting information

Suppelmental

## Conclusions

Few studies have assessed the phenotypic response of whole plants in terms of biomass allocation, organ morphology and anatomy simultaneously, and even less under multiple resource limitation. Here, our research demonstrates that major traits from all three categories shift in response to resource limitation. Many of these shifts are passive, i.e. scaling with decreased mass of plants under stressed conditions. Moreover, these passive shifts extend well beyond mass allocational traits and include both morphological and anatomical traits. The magnitude and direction of individual traits responses, and their coordination are driven not only by passive scaling with plant mass, but by an additional active component that can be complementary or antagonistic to the effect of mass scaling. Thus, for a full understanding of plants response to environmental stress both this passive and active plasticity needs to be taken into account.

## Acknowledgements

We would like to thank M. Boyd and K. Tarner for assistance with plant growth and data collection, K. Bettinger, S. Chhajed, G. Manning, J. Parrilli, N. Reisinger and J. Kobylanski for assistance with experimental setup and data collection, and the greater Donovan lab group for comments that improved this study and manuscript. Additionally, we would like to thank the reviewers for their time and constructive comments. This work was financially supported by grant NSF1444522 to LAD and a China Scholarship Council (CSC) grant to YW.

## Author contributions

YW, AT, and LD designed the experiment. YW carried out the experiment and took all measurements. YW and AT analyzed the results. YW wrote the initial manuscript draft with revisions by AT and LD. All authors contributed to manuscript revision, and read and approved the submitted version

---

**Table 1**. Trait response to a factorial of light and nutrient limitation. Light vs Shade designates direct sun vs 50% shade treatment. Rich vs Poor designates high nutrients vs 10% nutrient concentration treatment. For each trait, the ANOVA effects of genotype (G), light (L) and nutrient (N) stresses and their interactions (L×N) are presented, without (w/o) or with plant total biomass (Mass) included as a covariate: ***: p<0.001, **: p<0.01, *: p<0.05. Based on the difference between the models with and without biomass as covariate we assigned a treatment response as either passive (primarily scaling with mass), active (adjusted independent of mass), or a combination thereof. For each treatment combination, the mean±SE of 10 genotypes (estimated marginal mean from the ANOVA, based on 5-6 replicates per genotype) is presented for each trait. Different lower case letters represent significant (p<0.05) Tukey post hoc differences among treatments for each trait from the ANOVA without biomass as a covariate.

## Supplemental materials

**Supplemental Table S1.** List of genotypes used in this study, it’s common name, the corresponding plant ID from the USDA GRIN Database for each genotype, and the market type of each genotype.

**Supplemental Table S2.** Genotype means for all traits discussed

**Supplemental Figure S1**. Comparison of light levels between shaded and unshaded treatment. On average the low-light treatment received 50% of the photon flux density of the high-light treatment. Light intensities in each treatment were measured with a handheld light meter (LI-189; LI-COR, Lincoln, NE). Readings were taken from 8:00 AM to 7:00 PM. The light sensor was held near the soil level in each plot. We presented representative data from May 15th, a cloud-free day.

**Supplemental Figure S2**. Effect of accounting for size scaling in traits when calculating relative distance plasticity index (RDPI). RDPI changes are shown per treatment (Light/Shade, full sun/50% shade; Rich/Poor, full nutrients/10% nutrients) and whether RDPI measures were positive or negative when initially compared to control (Light-Rich). For positive RDPI values (traits that increase in trait value with stress) a positive change when taking size scaling into account shows that size scaling decreased apparent (as measured initially) plasticity (i.e taking size scaling into account shows increased plasticity). A negative change on the other hand shows that the apparent plasticity was enhanced by size scaling (i.e. taking size scaling into account shows decreased plasticity. For negative RDPI values (traits that decrease in trait value with stress) the effect of the sign of change is reversed. Symbols indicate whether RDPI significance (different from zero) is gained (circles), lost (squares), or remains unchanged (triangles, when taking biomass into account.

**Supplemental Figure S3.** Scaling relationship between biomass and trait value among and between genotypes and treatments. **(a)** Scaling relationships between individual plants per genotypes and treatment (dotted lines) and **(b)** scaling relationships between genotypes per treatment.

**Supplemental Figure S4.** Trait contributions to principal components of figure 5.

## Notes

### Competing Interest Statement

The authors have declared no competing interest.

### Summary of Updates

Thanks to reviewer feedback updated the direction and focus of the paper towards allometric scaling and plasticity due to resource limitation.

