## Supplementary material for "Plasticity and the role of mass-scaling in allocation, morphology and anatomical trait responses to above and belowground resource limitation in cultivated sunflower (*Helianthus annuus L.*)": Suppelmental

**Table S1.** List of genotypes used in this study, it's common name, the corresponding plant ID from the USDA GRIN Database for each genotype, and the market type of each genotype.

| Genotype ID | USDA GRIN ID | Common name | Market Type |
|---|---|---|---|
| *8* | PI 552943 | RHA 280 | RHA-NonOil |
| *11* | PI 561920 | HA 380 | HA-NonOil |
| *12* | PI 561921 | RHA 381 | RHA-Oil |
| *16* | PI 578011 | RHA 389 | RHA-Oil |
| *31* | PI 607921 | R-188 | RHA-Oil |
| *98* | PI 664201 | RHA 326 | RHA-NonOil |
| *126* | PI 549006 | HA GERMPLASM POOL III-L | Oil introgressed |
| *227* | PI 650753 | HA-R2 | HA-Oil |
| *231* | PI 509051 | HA 341 | HA-Oil |
| *252* | PI 618727 | HA 423 | HA-Oil |

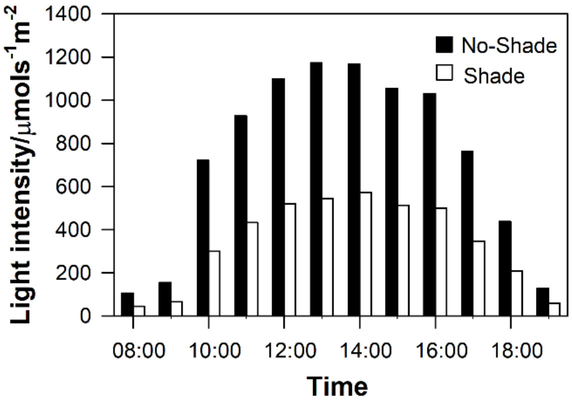


**Fig S1**. Comparison of light levels between shaded and unshaded treatment. On average the low-light treatment received 50% of the photon flux density of the high-light treatment. Light intensities in each treatment were measured with a handheld light meter (LI-189; LI-COR, Lincoln, NE). Readings were taken from 8:00 AM to 7:00 PM. The light sensor was held near the sand level in each plot. We presented representative data from May 15th, a cloud-free day.

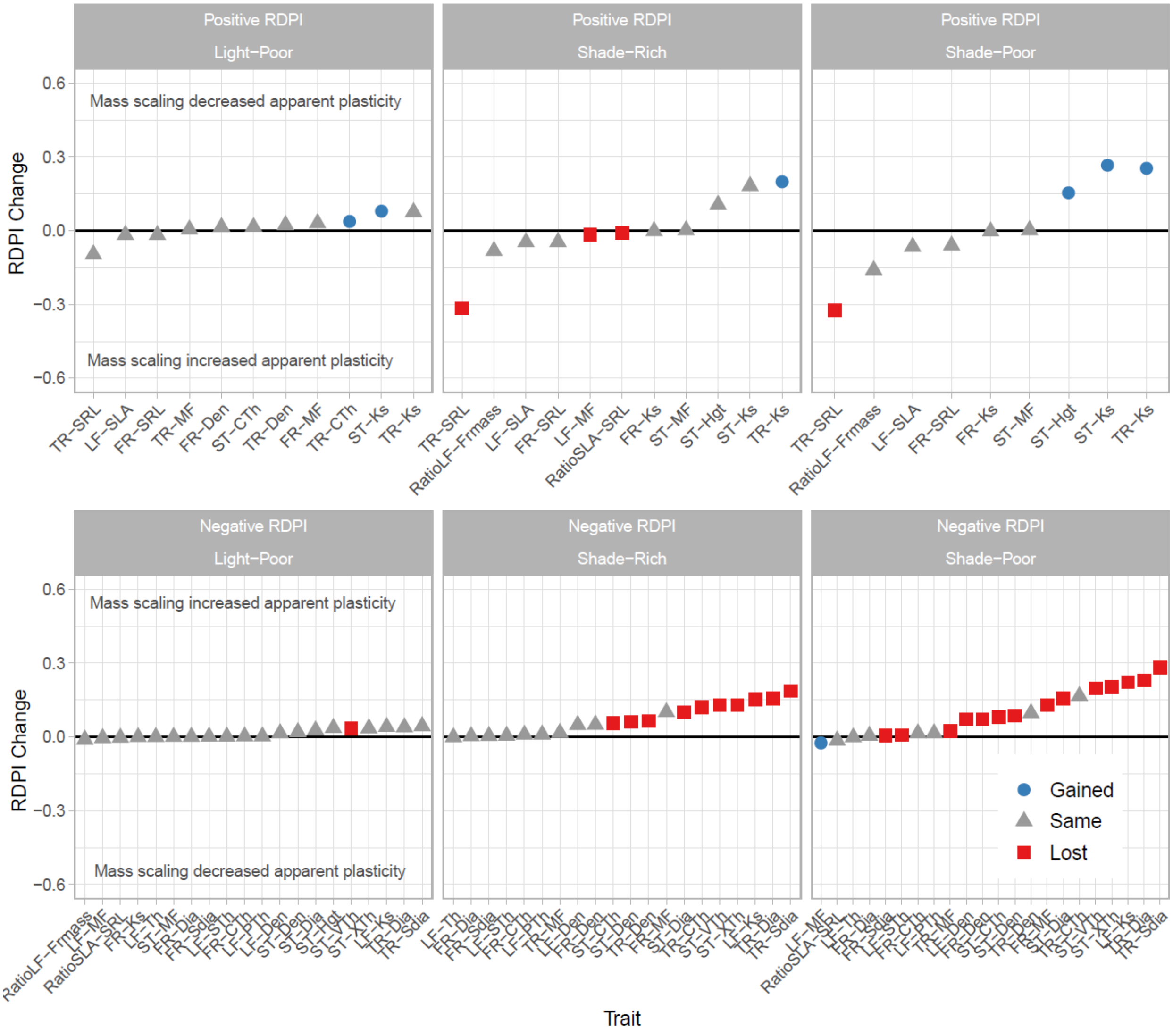


**Fig S2**. Effect of accounting for mass scaling in traits when calculating relative distance plasticity index (RDPI). RDPI changes are shown per treatment (Light/Shade, full sun/50% shade; Rich/Poor, full nutrients/10% nutrients) and whether RDPI measures were positive or negative when initially compared to control (Light-Rich). For positive RDPI values (traits that increase in trait value with stress) a positive change when taking mass scaling into account shows that mass scaling decreased apparent (as measured initially) plasticity (i.e taking size scaling into account shows increased plasticity). A negative change on the other hand shows that the apparent plasticity was enhanced by mass scaling (i.e taking size scaling into account shows decreased plasticity. For negative RDPI values (traits that decrease in trait value with stress) the effect of the sign of change is reversed. Symbols indicate whether RDPI significance (different from zero) is gained (circles), lost (squares), or remains unchanged (triangles, when taking biomass into account.

**Fig S3.** Mass scaling relationship between biomass and trait value within genotypes and across genotypes per treatments. (a) Mass scaling relationships per genotype (dotted lines) based on individual plants (points) with treatment fit line (solid lines)  (b) Mass scaling relationship across genotypes (points) per treatment (solid lines) .

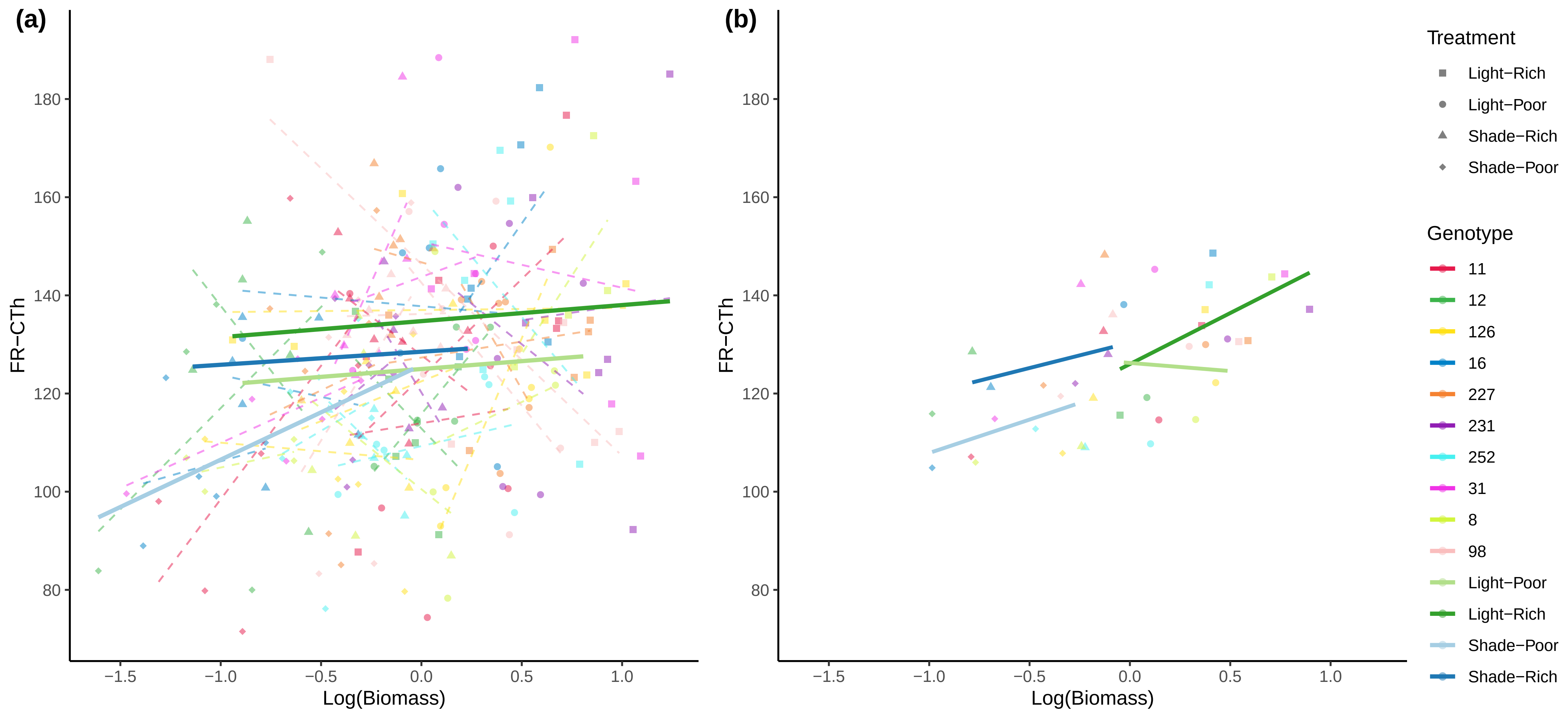

(a)
(b)
FR−CTh
Log(Biomass)
180
160
140
120
100
80
−1.5
−1.0
−0.5
0.0
0.5
1.0
Treatment
Light−Rich
Light−Poor
Shade−Rich
Shade−Poor
Genotype
11
12
126
16
227
231
252
31
8
98
Light−Poor
Light−Rich
Shade−Poor
Shade−Rich

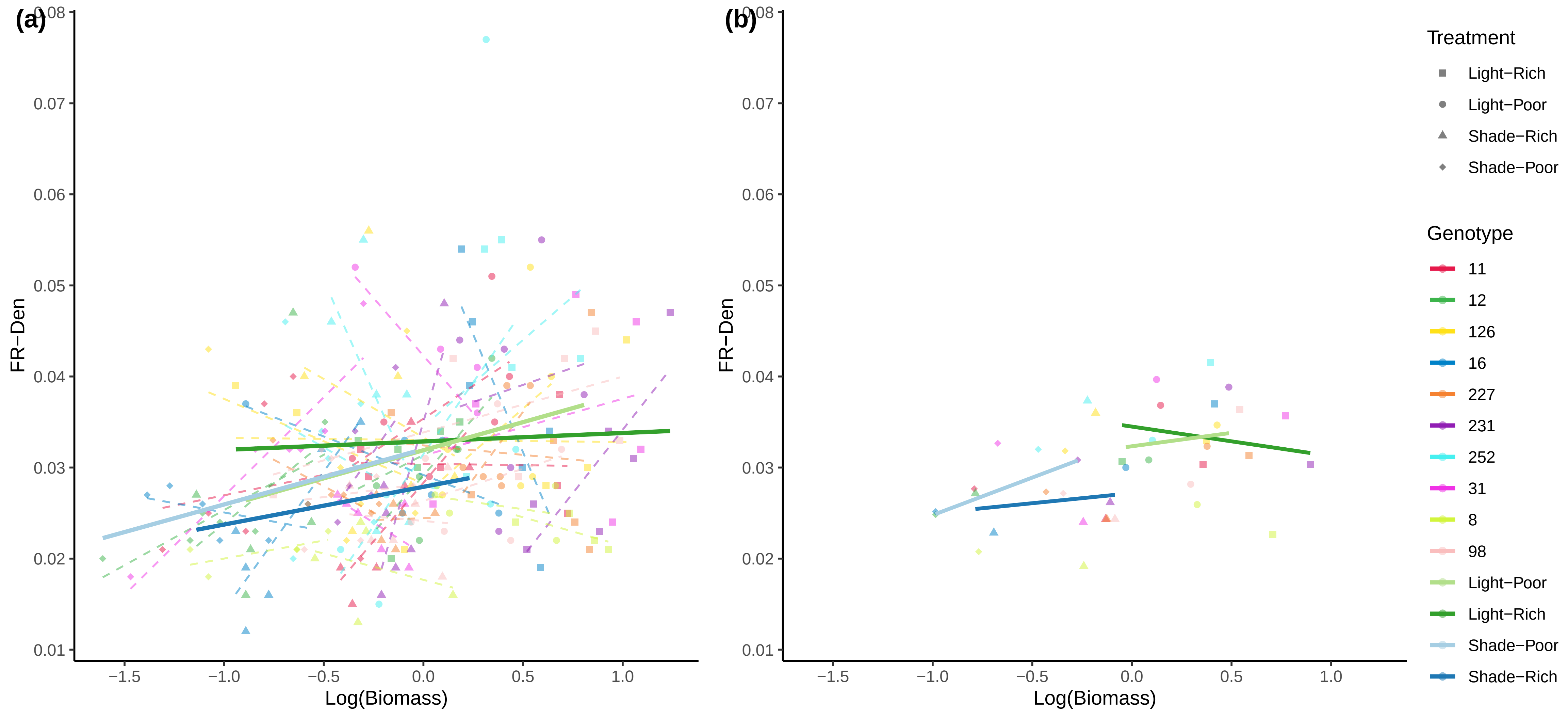
(a)
(b)
FR−Den
Log(Biomass)
0.08
0.07
0.06
0.05
0.04
0.03
0.02
0.01
−1.5
−1.0
−0.5
0.0
0.5
1.0
Treatment
Light−Rich
Light−Poor
Shade−Rich
Shade−Poor
Genotype
11
12
126
16
227
231
252
31
8
98
Light−Poor
Light−Rich
Shade−Poor
Shade−Rich

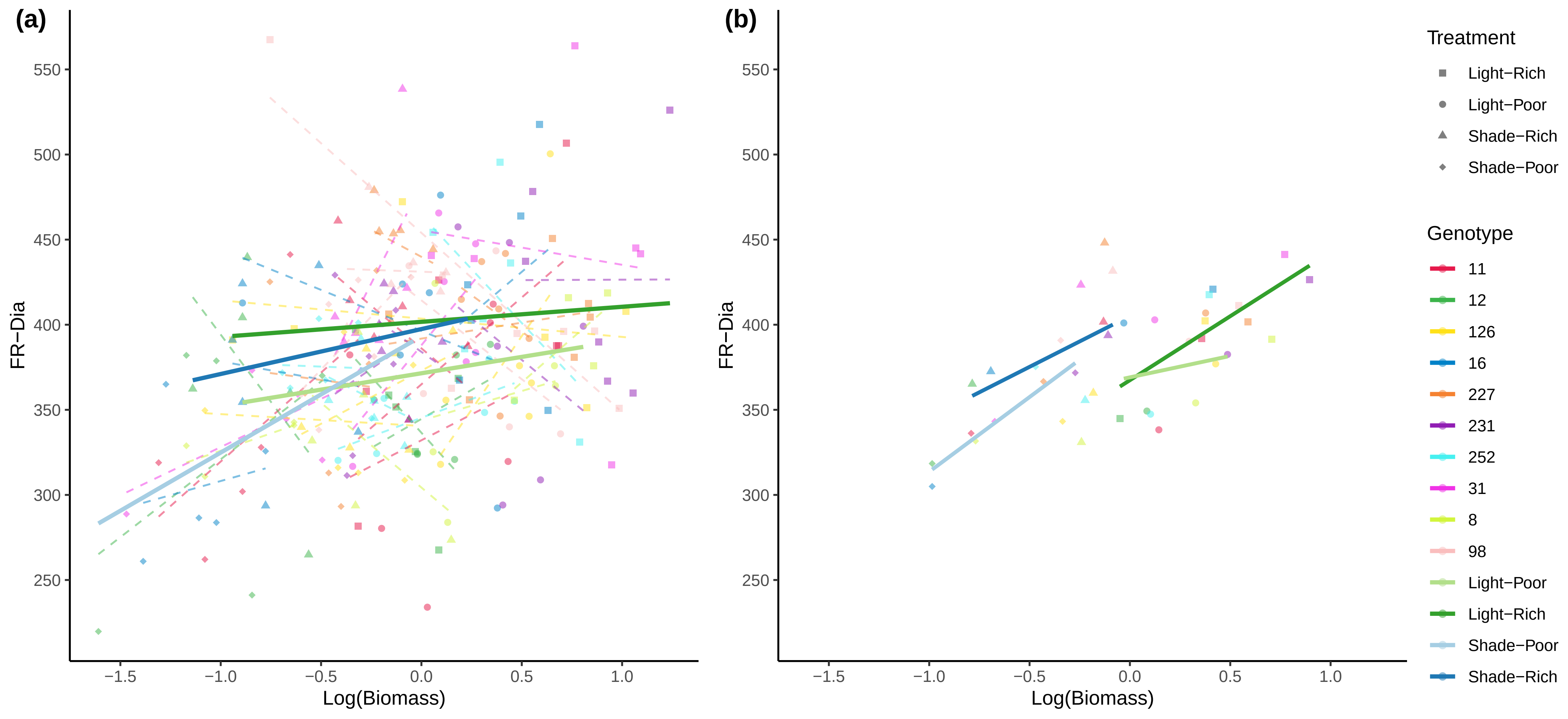

(a)
(b)
FR−Dia
Log(Biomass)
550
500
450
400
350
300
250
−1.5
−1.0
−0.5
0.0
0.5
1.0
Treatment
Light−Rich
Light−Poor
Shade−Rich
Shade−Poor
Genotype
11
12
126
16
227
231
252
31
8
98
Light−Poor
Light−Rich
Shade−Poor
Shade−Rich

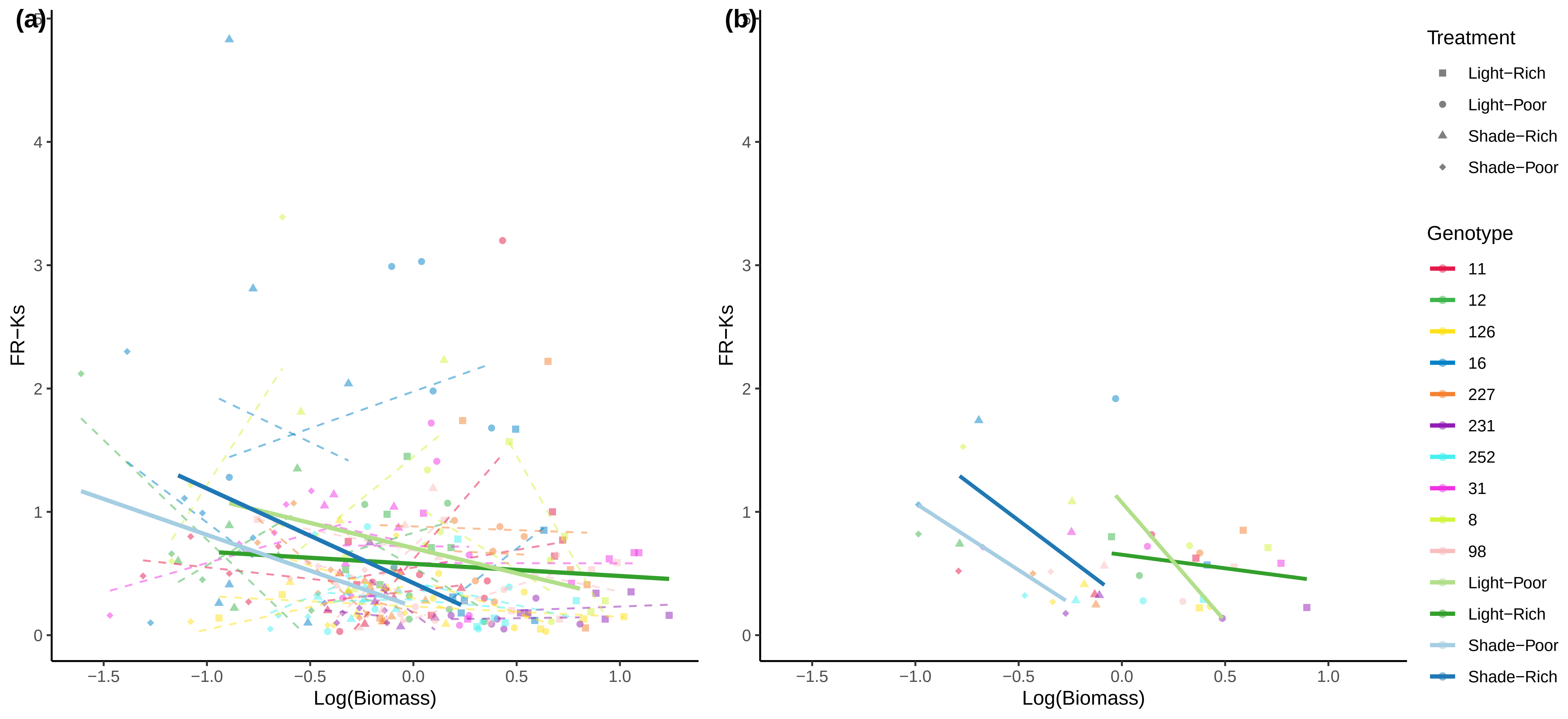

(a)
(b)
FR−Ks
Log(Biomass)
Treatment
Light−Rich
Light−Poor
Shade−Rich
Shade−Poor
Genotype
11
12
126
16
227
231
252
31
8
98
Light−Poor
Light−Rich
Shade−Poor
Shade−Rich

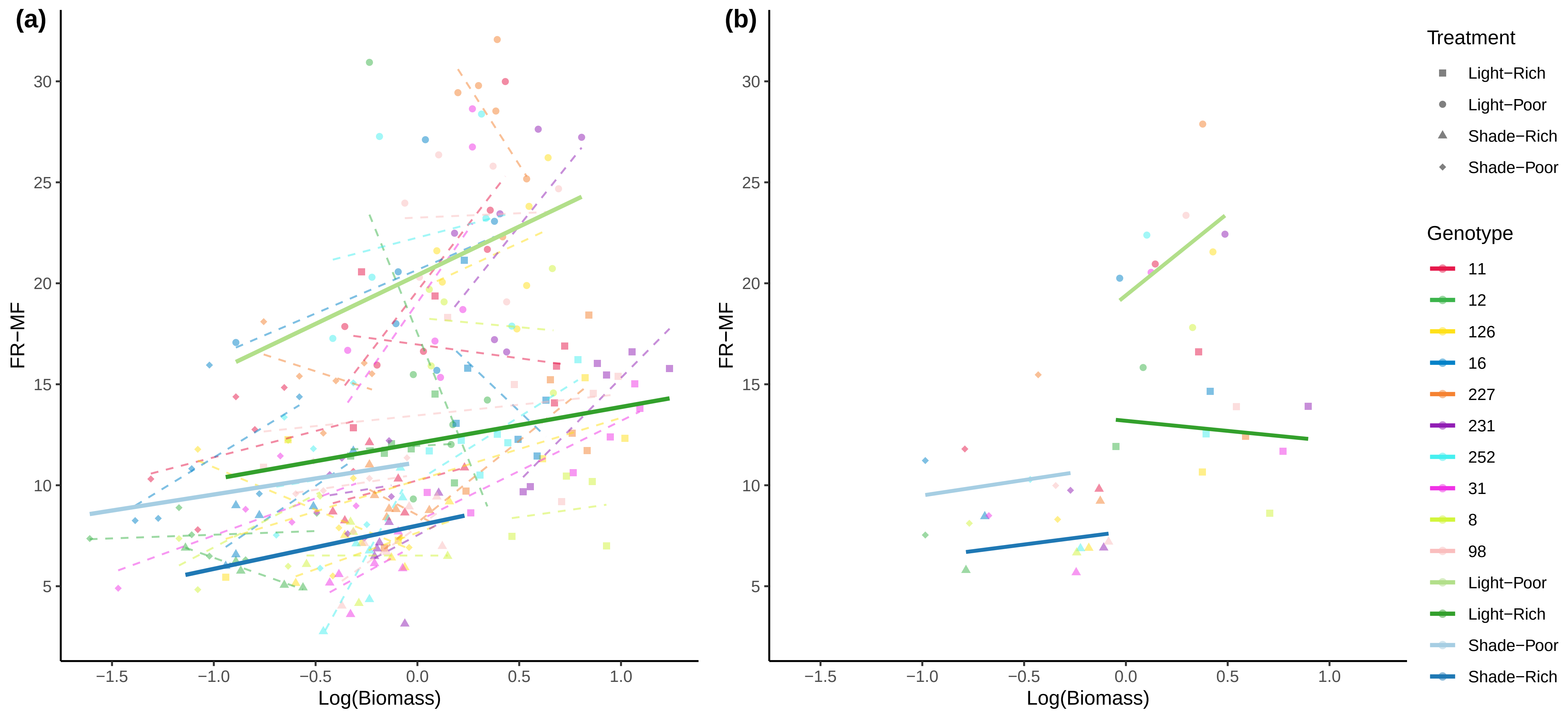

(a)
(b)
FR−MF
Log(Biomass)
Treatment
Light−Rich
Light−Poor
Shade−Rich
Shade−Poor
Genotype
11
12
126
16
227
231
252
31
8
98
Light−Poor
Light−Rich
Shade−Poor
Shade−Rich

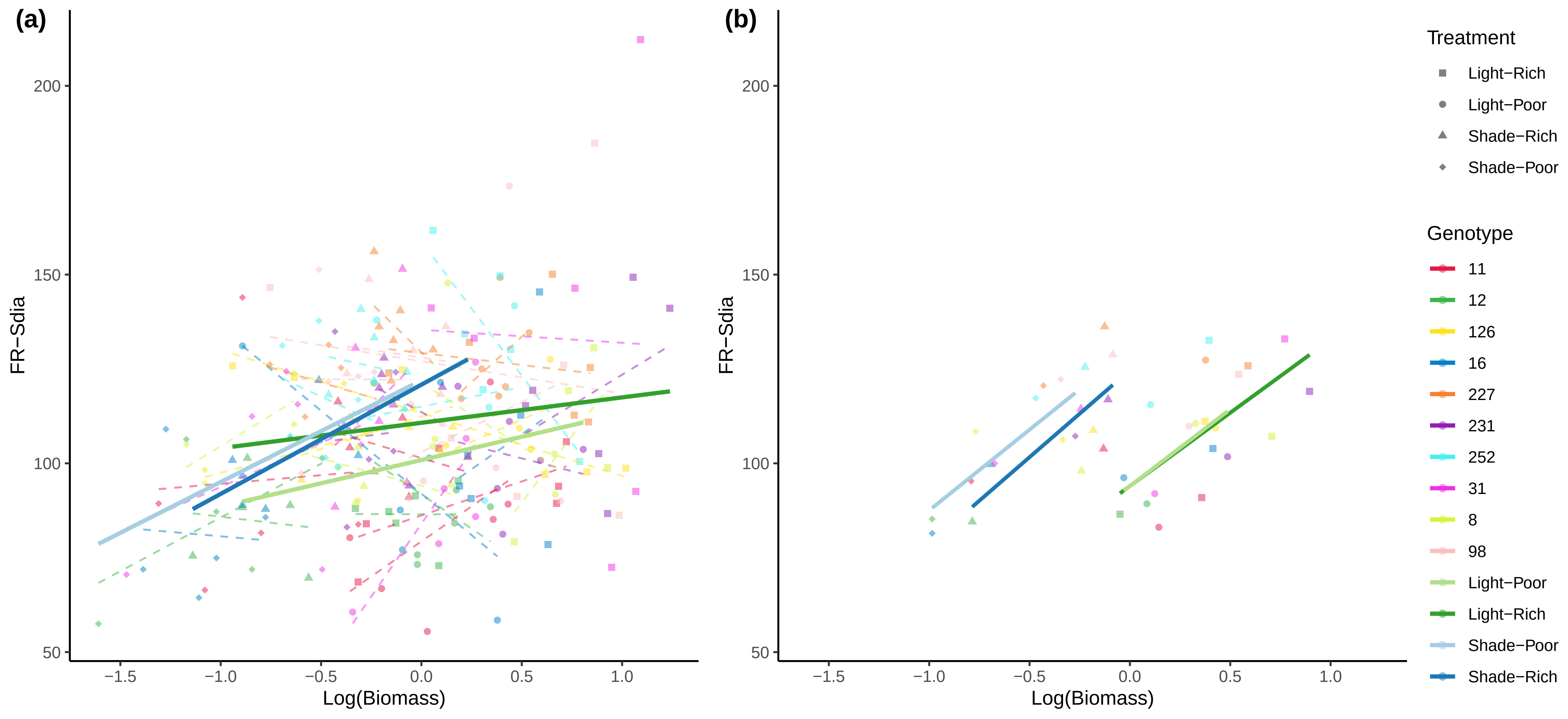

(a)
(b)
FR−Sdia
Log(Biomass)
200
150
100
50
−1.5
−1.0
−0.5
0.0
0.5
1.0
Treatment
Light−Rich
Light−Poor
Shade−Rich
Shade−Poor
Genotype
11
12
126
16
227
231
252
31
8
98
Light−Poor
Light−Rich
Shade−Poor
Shade−Rich

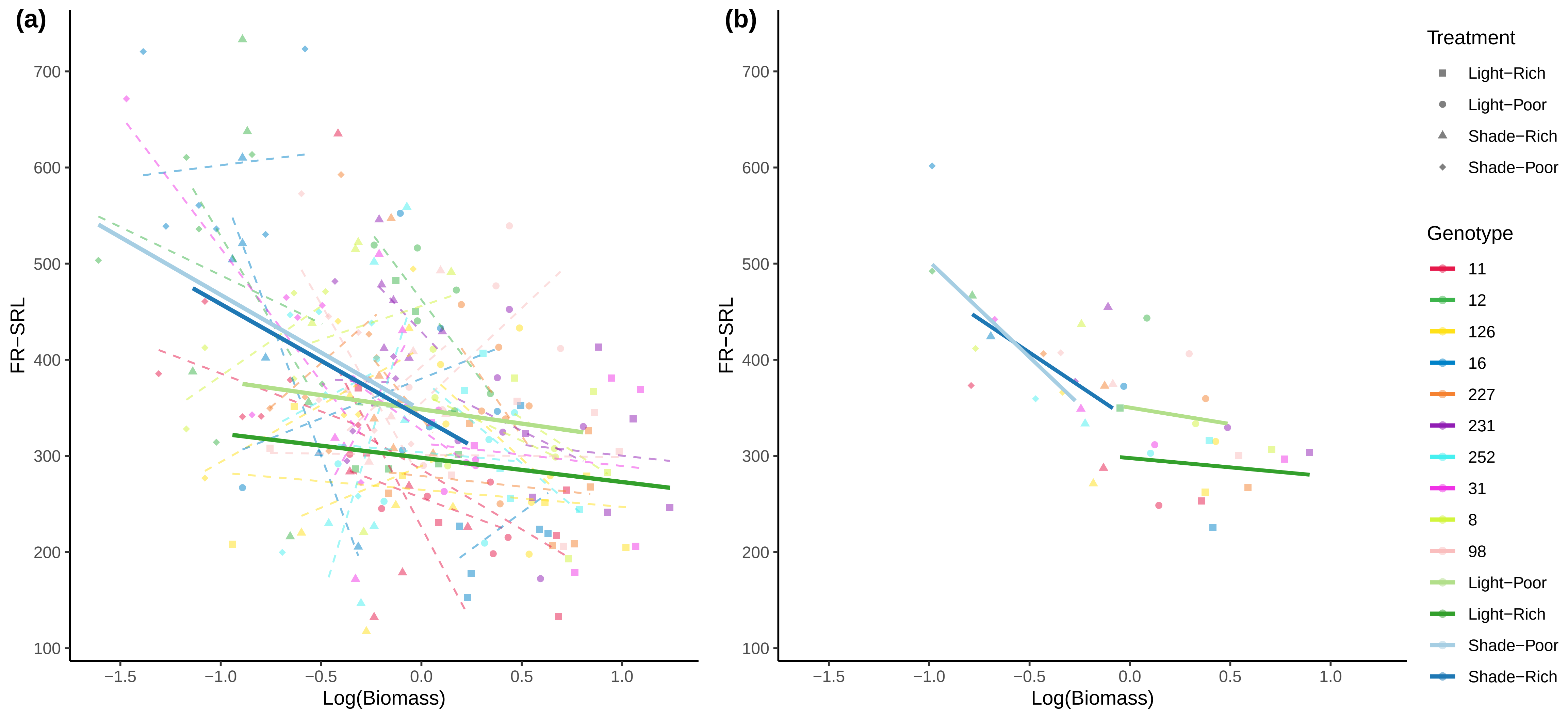

(a)
(b)
FR−SRL
Log(Biomass)
700
600
500
400
300
200
100
−1.5
−1.0
−0.5
0.0
0.5
1.0
Treatment
Light−Rich
Light−Poor
Shade−Rich
Shade−Poor
Genotype
11
12
126
16
227
231
252
31
8
98
Light−Poor
Light−Rich
Shade−Poor
Shade−Rich

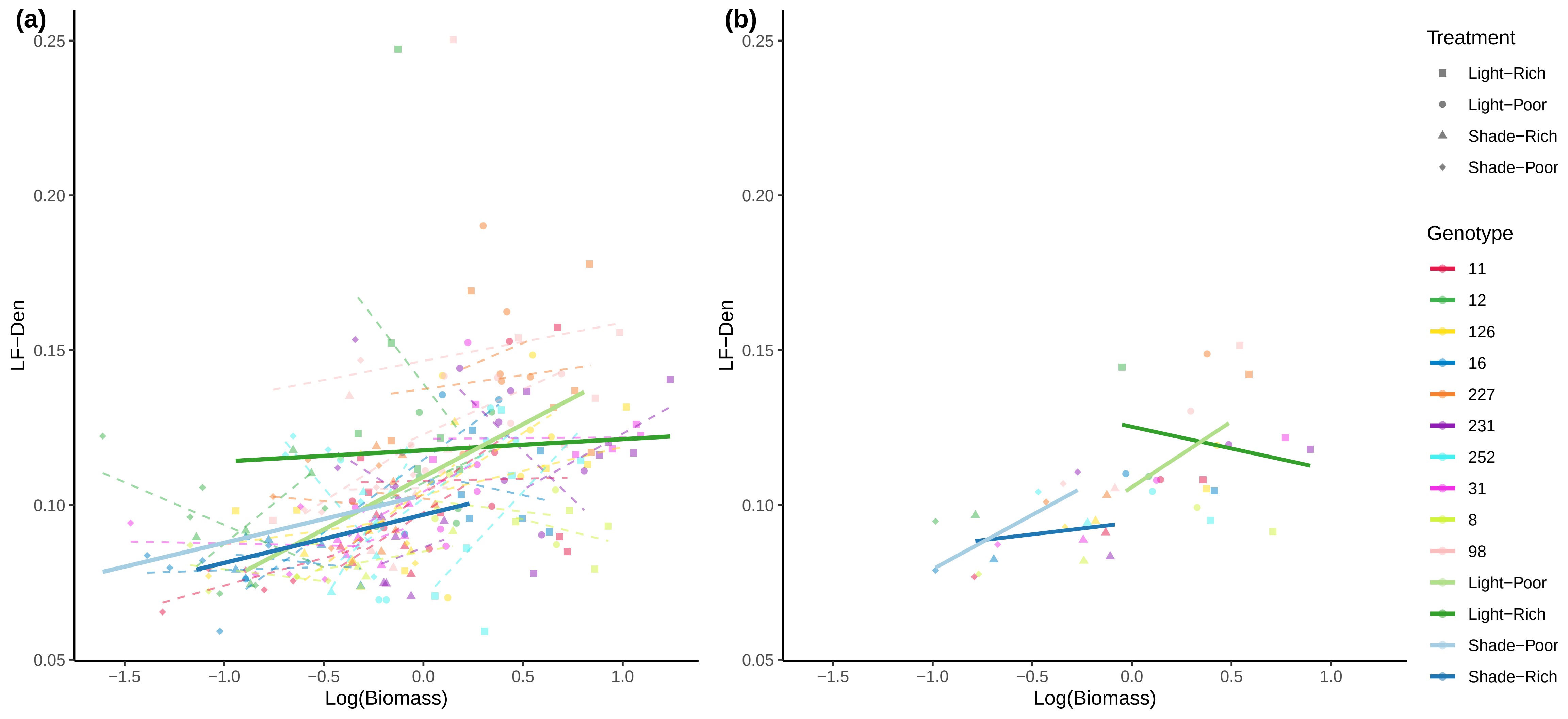

(a)
(b)
LF−Den
Log(Biomass)
0.25
0.20
0.15
0.10
0.05
−1.5
−1.0
−0.5
0.0
0.5
1.0
Treatment
Light−Rich
Light−Poor
Shade−Rich
Shade−Poor
Genotype
11
12
126
16
227
231
252
31
8
98
Light−Poor
Light−Rich
Shade−Poor
Shade−Rich

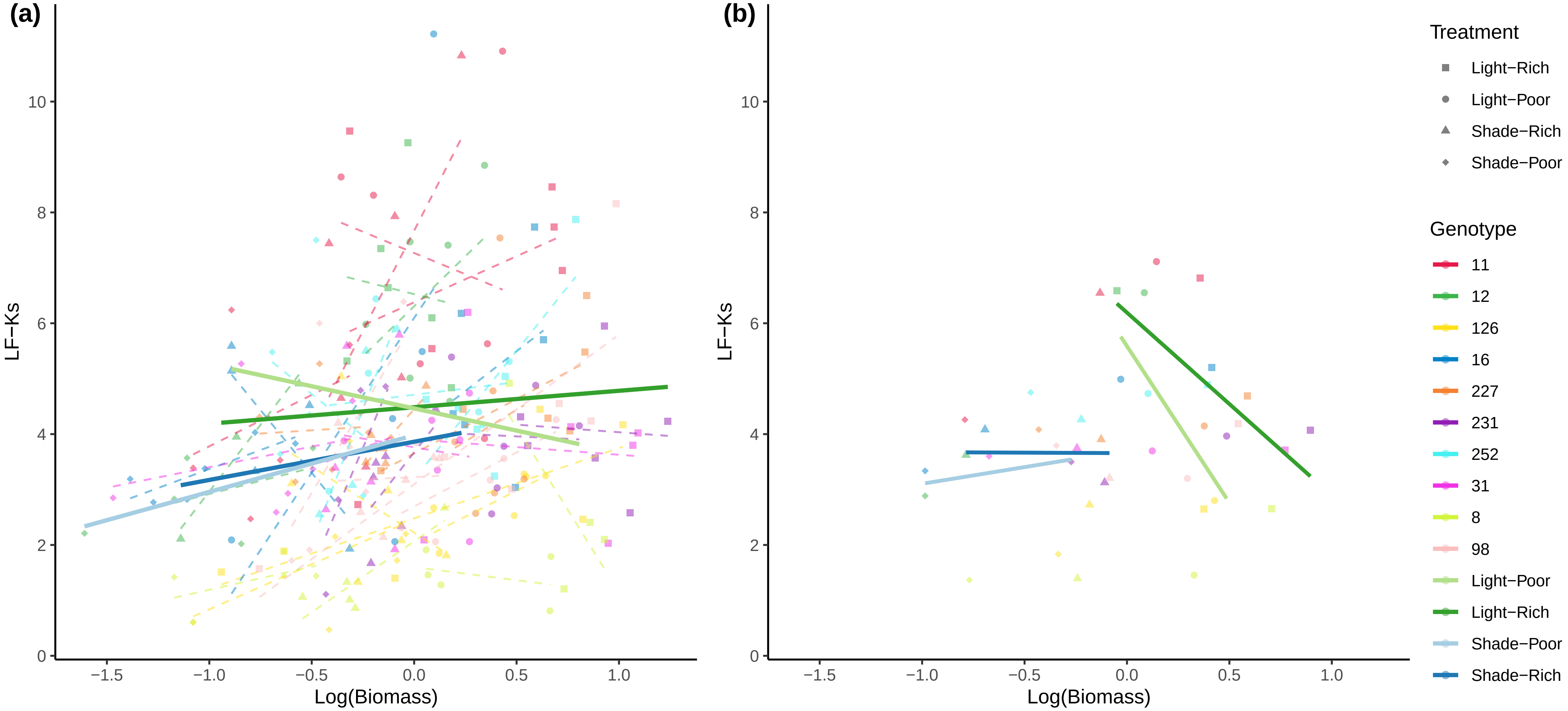

(a)
(b)
LF−Ks
Log(Biomass)
Treatment
Light−Rich
Light−Poor
Shade−Rich
Shade−Poor
Genotype
11
12
126
16
227
231
252
31
8
98
Light−Poor
Light−Rich
Shade−Poor
Shade−Rich

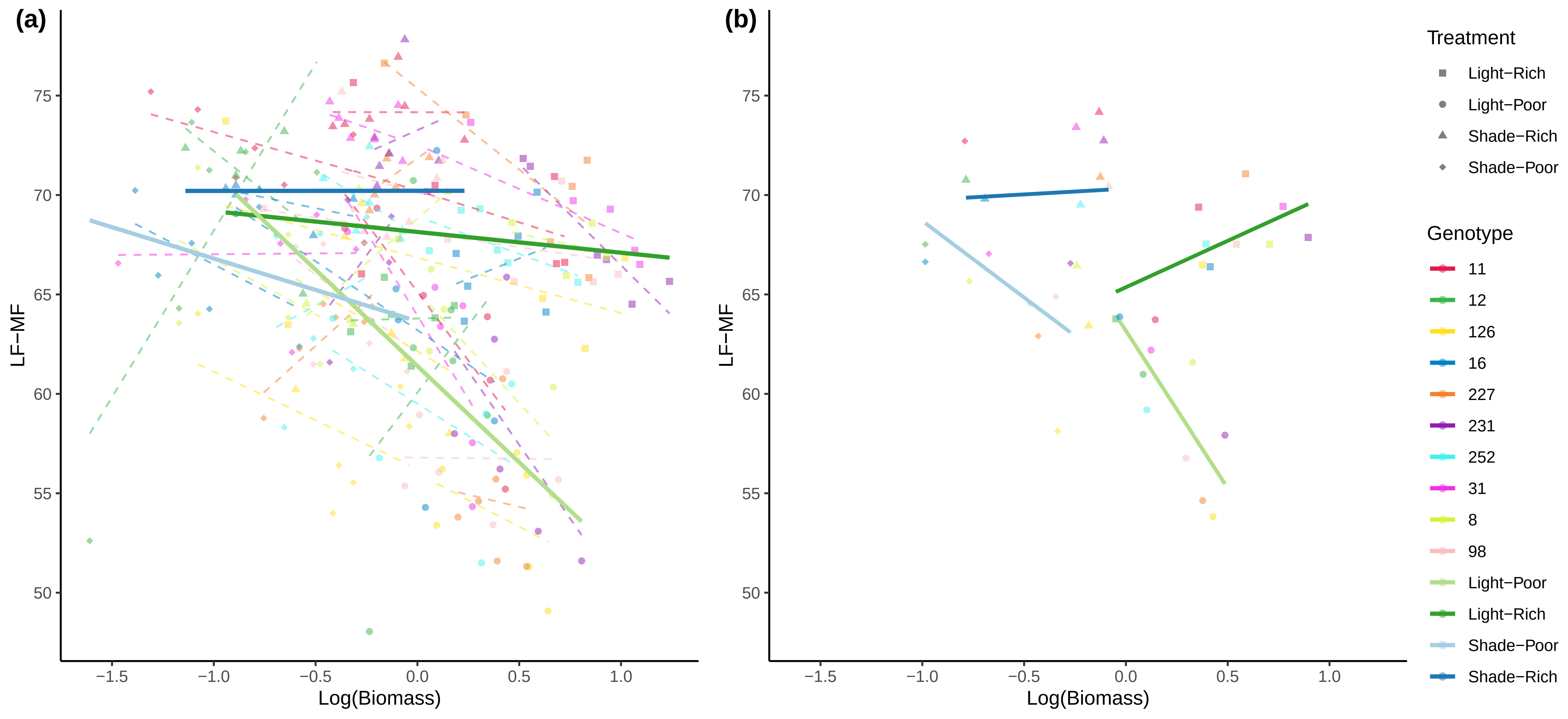

(a)
(b)
LF−MF
Log(Biomass)
75
70
65
60
55
50
−1.5
−1.0
−0.5
0.0
0.5
1.0
Treatment
Light−Rich
Light−Poor
Shade−Rich
Shade−Poor
Genotype
11
12
126
16
227
231
252
31
8
98
Light−Poor
Light−Rich
Shade−Poor
Shade−Rich

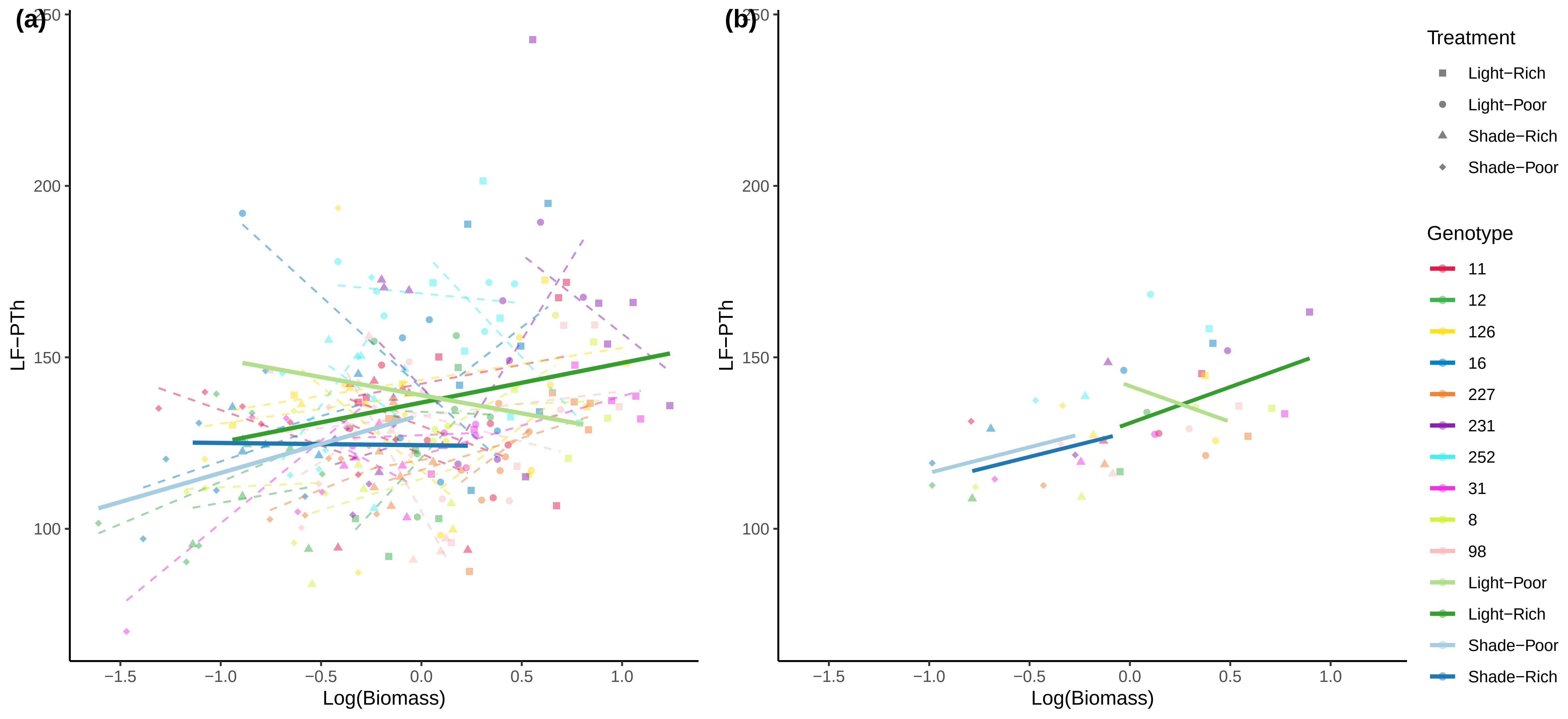

(a)
(b)
250
200
150
100
LF−PTh
−1.5
−1.0
−0.5
0.0
0.5
1.0
Log(Biomass)
Treatment
Light−Rich
Light−Poor
Shade−Rich
Shade−Poor
Genotype
11
12
126
16
227
231
252
31
8
98
Light−Poor
Light−Rich
Shade−Poor
Shade−Rich

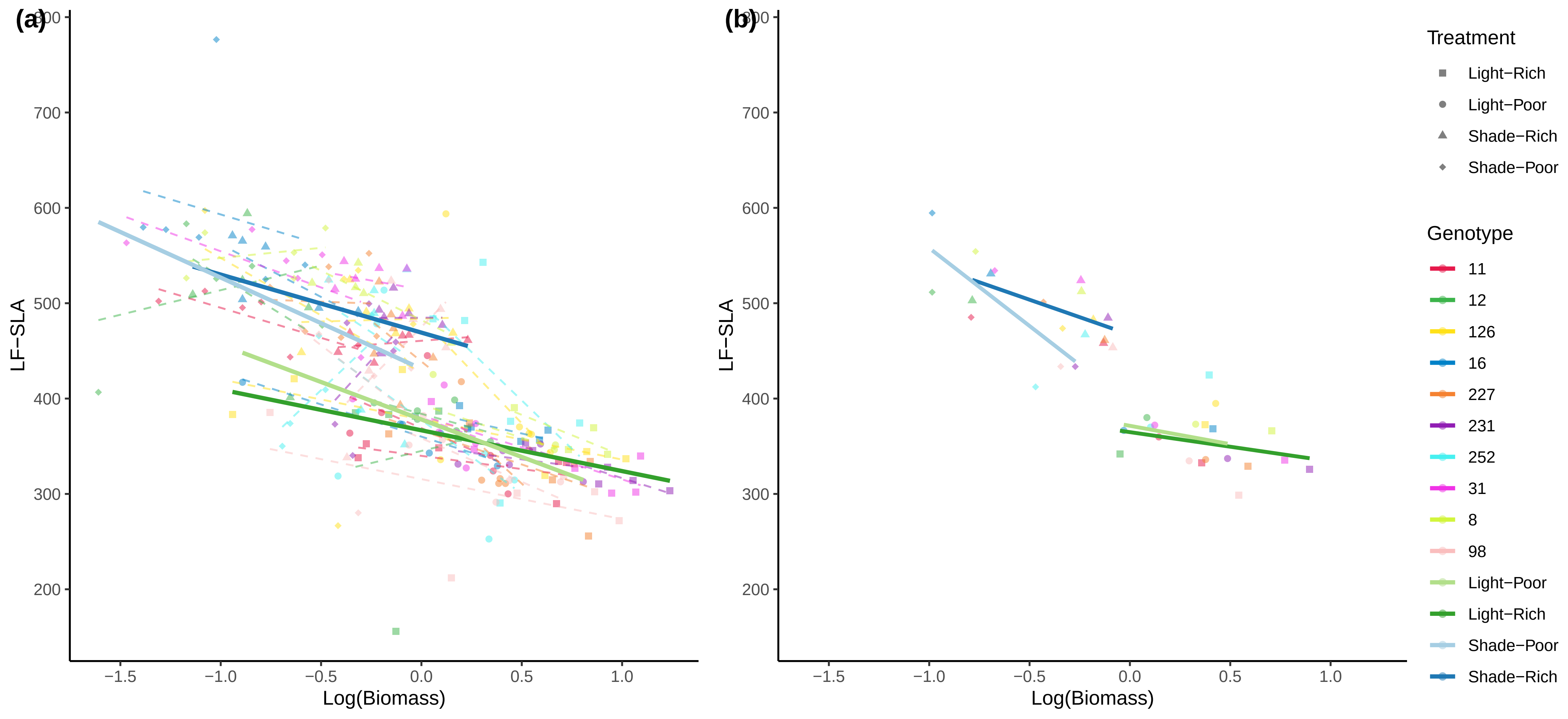

(a)
(b)
800
700
600
500
400
300
200
LF−SLA
−1.5
−1.0
−0.5
0.0
0.5
1.0
Log(Biomass)
Treatment
Light−Rich
Light−Poor
Shade−Rich
Shade−Poor
Genotype
11
12
126
16
227
231
252
31
8
98
Light−Poor
Light−Rich
Shade−Poor
Shade−Rich

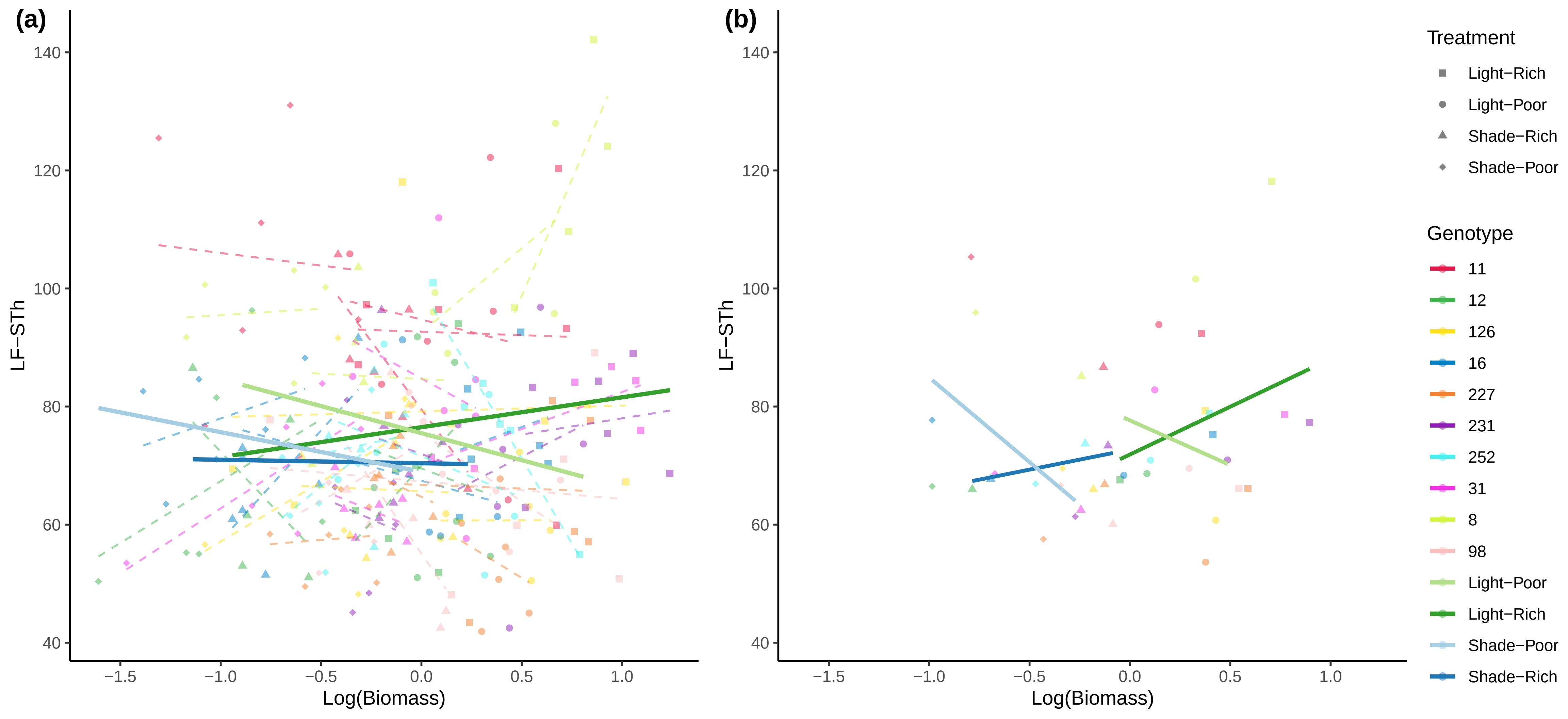

(a)
(b)
LF−STh
Log(Biomass)
140
120
100
80
60
40
−1.5
−1.0
−0.5
0.0
0.5
1.0
Treatment
Light−Rich
Light−Poor
Shade−Rich
Shade−Poor
Genotype
11
12
126
16
227
231
252
31
8
98
Light−Poor
Light−Rich
Shade−Poor
Shade−Rich

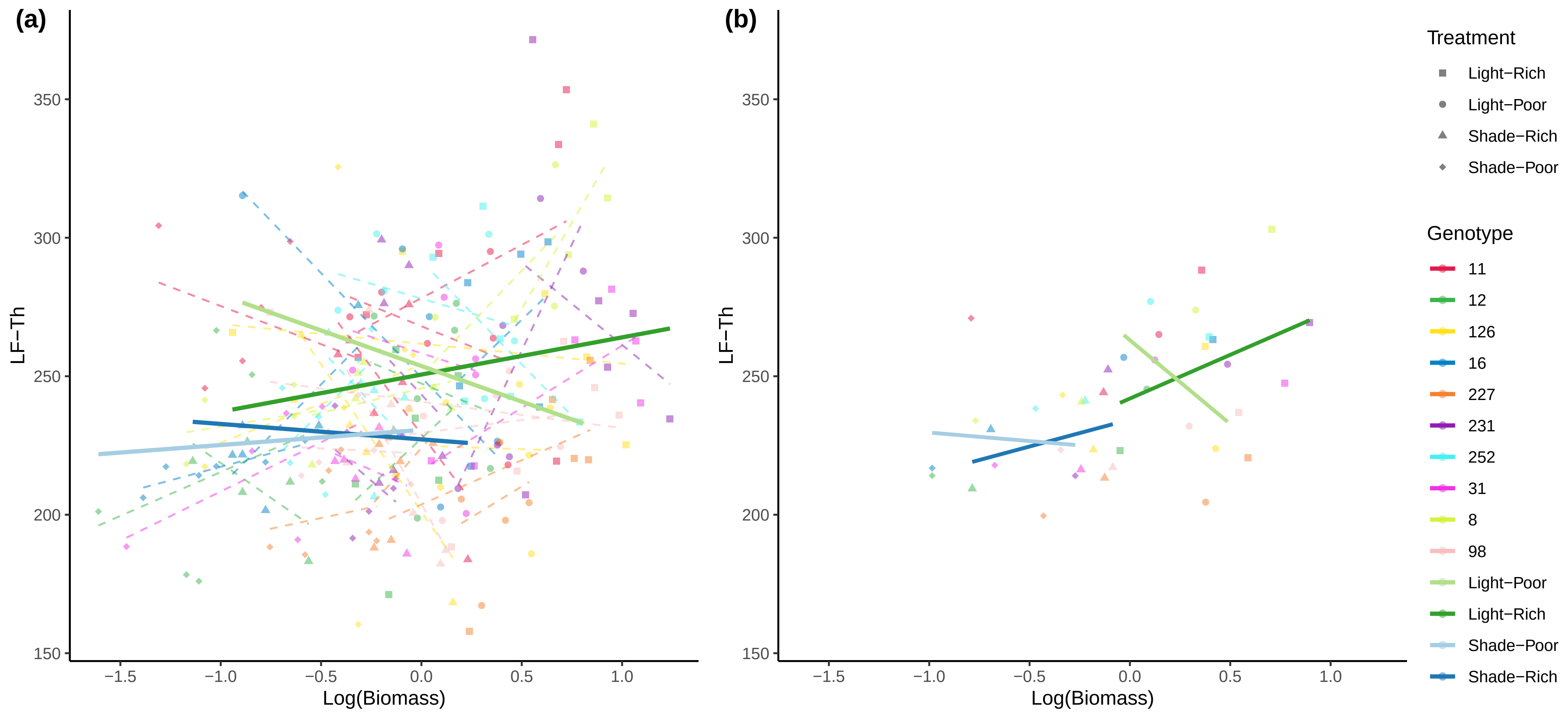

(a)
(b)
LF−Th
Log(Biomass)
150
200
250
300
350
−1.5
−1.0
−0.5
0.0
0.5
1.0
Treatment
Light−Rich
Light−Poor
Shade−Rich
Shade−Poor
Genotype
11
12
126
16
227
231
252
31
8
98
Light−Poor
Light−Rich
Shade−Poor
Shade−Rich

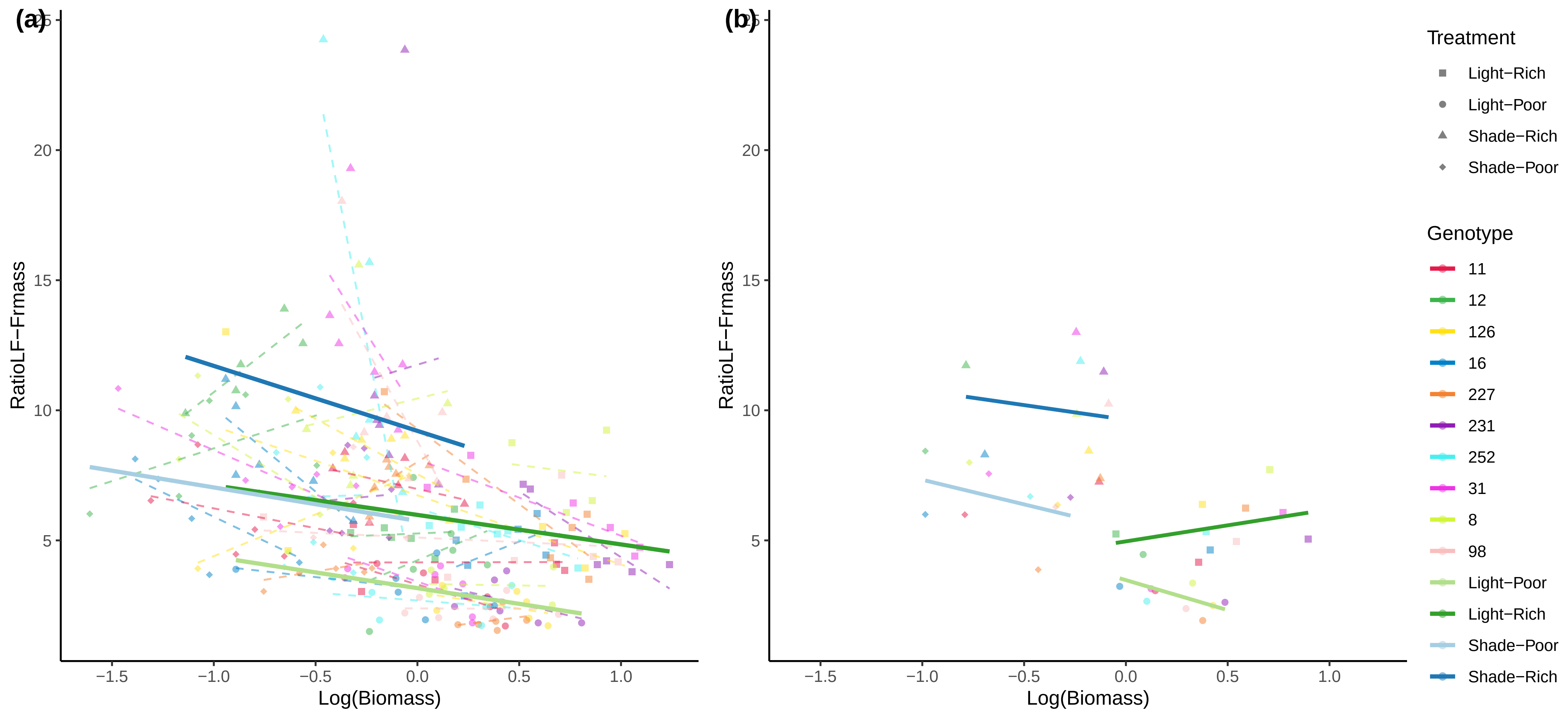

(a)
(b)
RatioLF−Frmass
Log(Biomass)
25
20
15
10
5
−1.5
−1.0
−0.5
0.0
0.5
1.0
Treatment
Light−Rich
Light−Poor
Shade−Rich
Shade−Poor
Genotype
11
12
126
16
227
231
252
31
8
98
Light−Poor
Light−Rich
Shade−Poor
Shade−Rich

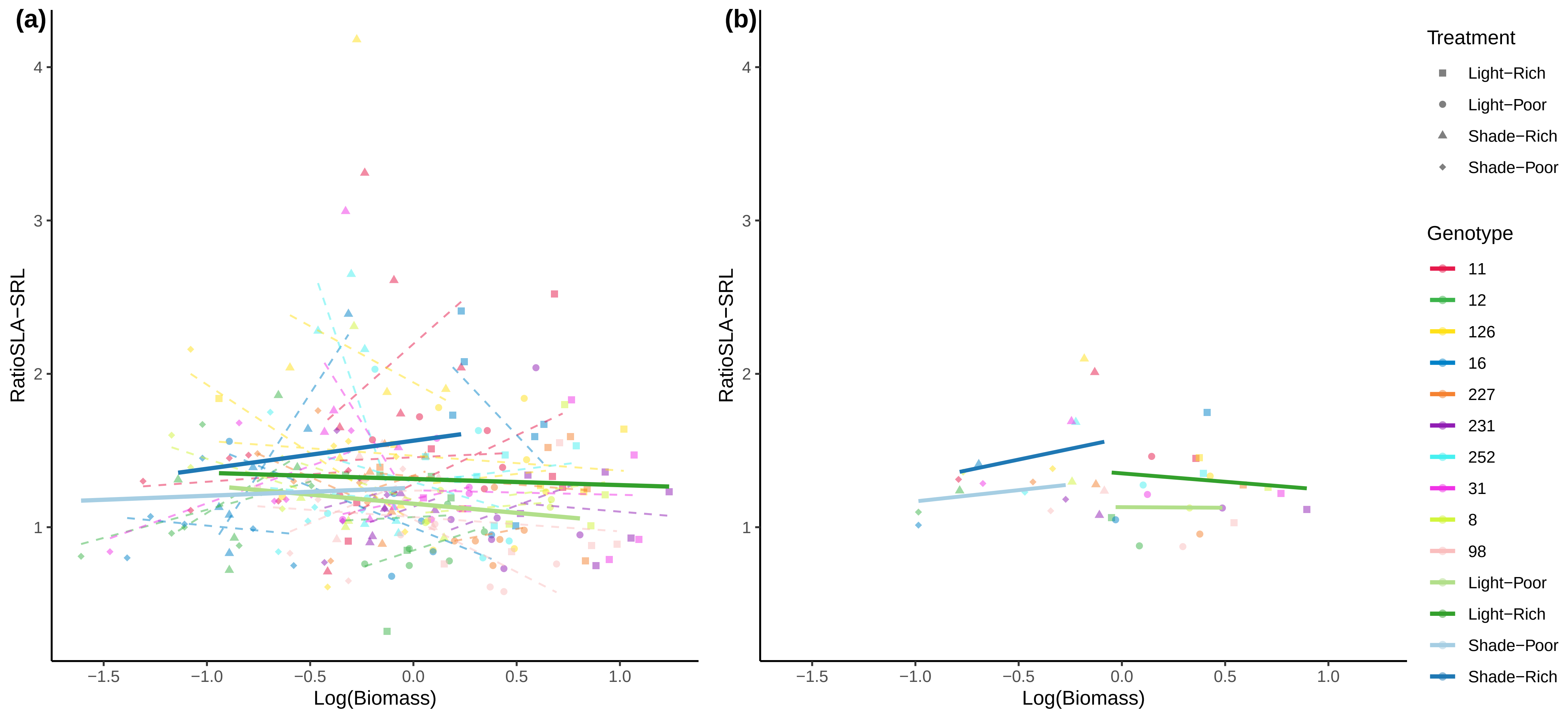

(a)
(b)
RatioSLA−SRL
Log(Biomass)
−1.5
−1.0
−0.5
0.0
0.5
1.0
1
2
3
4
Treatment
Light−Rich
Light−Poor
Shade−Rich
Shade−Poor
Genotype
11
12
126
16
227
231
252
31
8
98
Light−Poor
Light−Rich
Shade−Poor
Shade−Rich

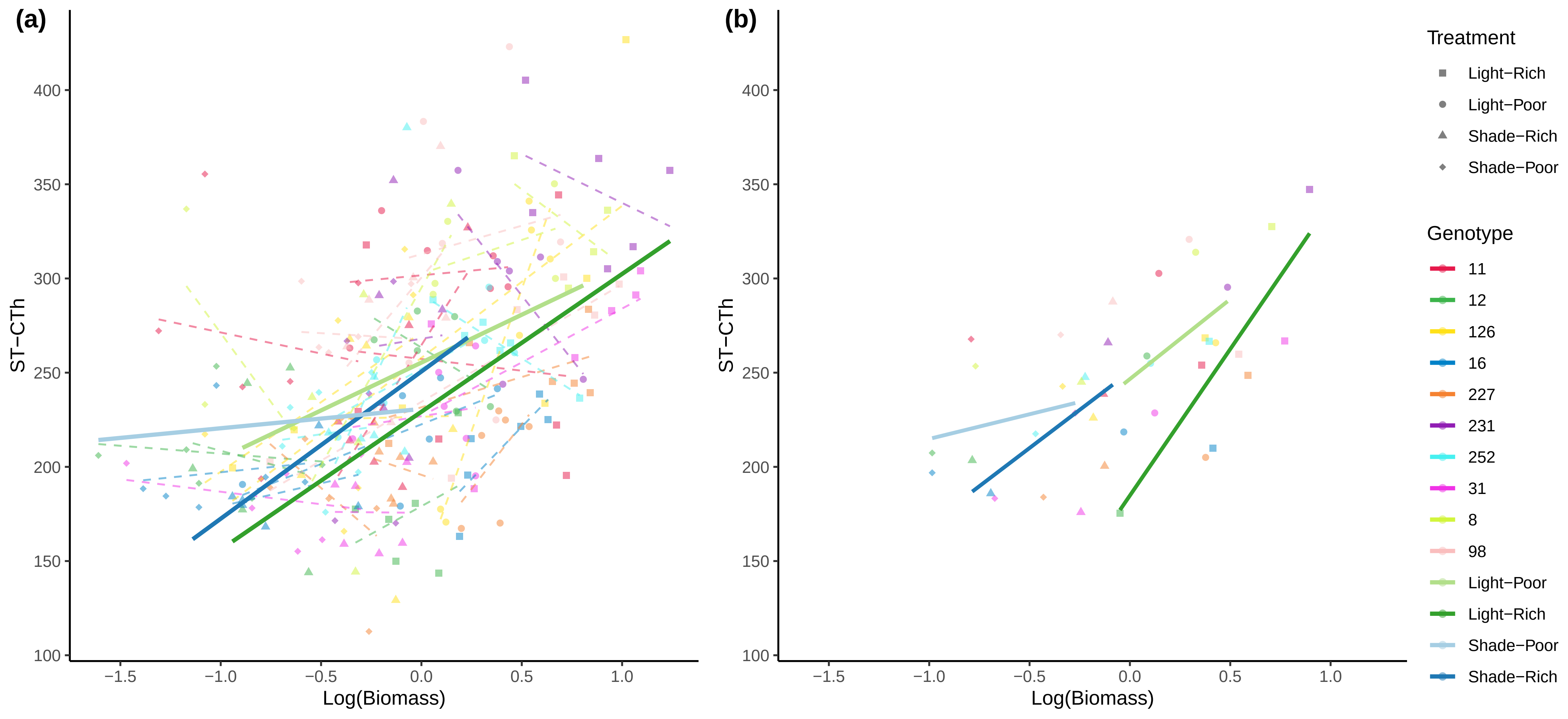

(a)
(b)
ST−CTh
Log(Biomass)
100
150
200
250
300
350
400
−1.5
−1.0
−0.5
0.0
0.5
1.0
Treatment
Light−Rich
Light−Poor
Shade−Rich
Shade−Poor
Genotype
11
12
126
16
227
231
252
31
8
98
Light−Poor
Light−Rich
Shade−Poor
Shade−Rich

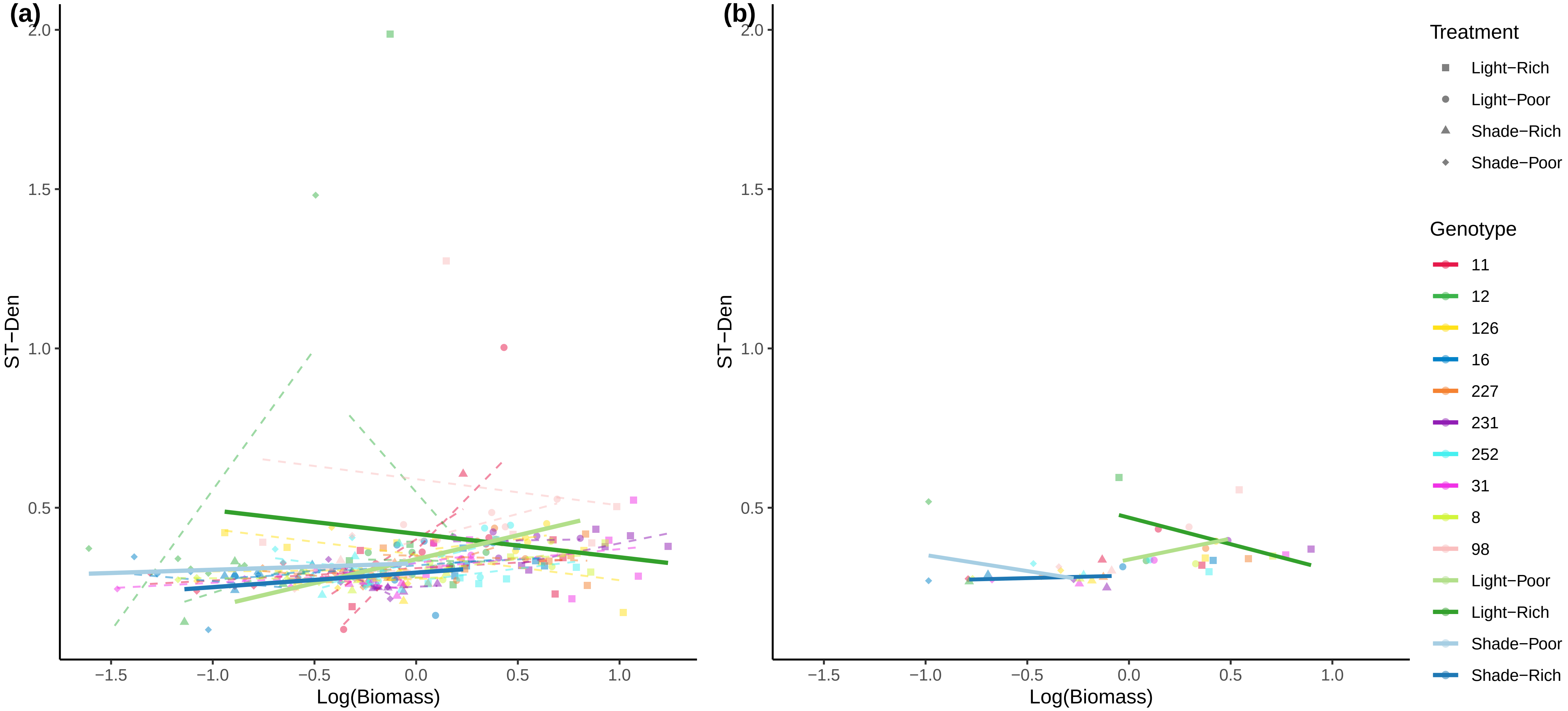
(a)
(b)
ST−Den
Log(Biomass)
2.0
1.5
1.0
0.5
−1.5
−1.0
−0.5
0.0
0.5
1.0
Treatment
Light−Rich
Light−Poor
Shade−Rich
Shade−Poor
Genotype
11
12
126
16
227
231
252
31
8
98
Light−Poor
Light−Rich
Shade−Poor
Shade−Rich

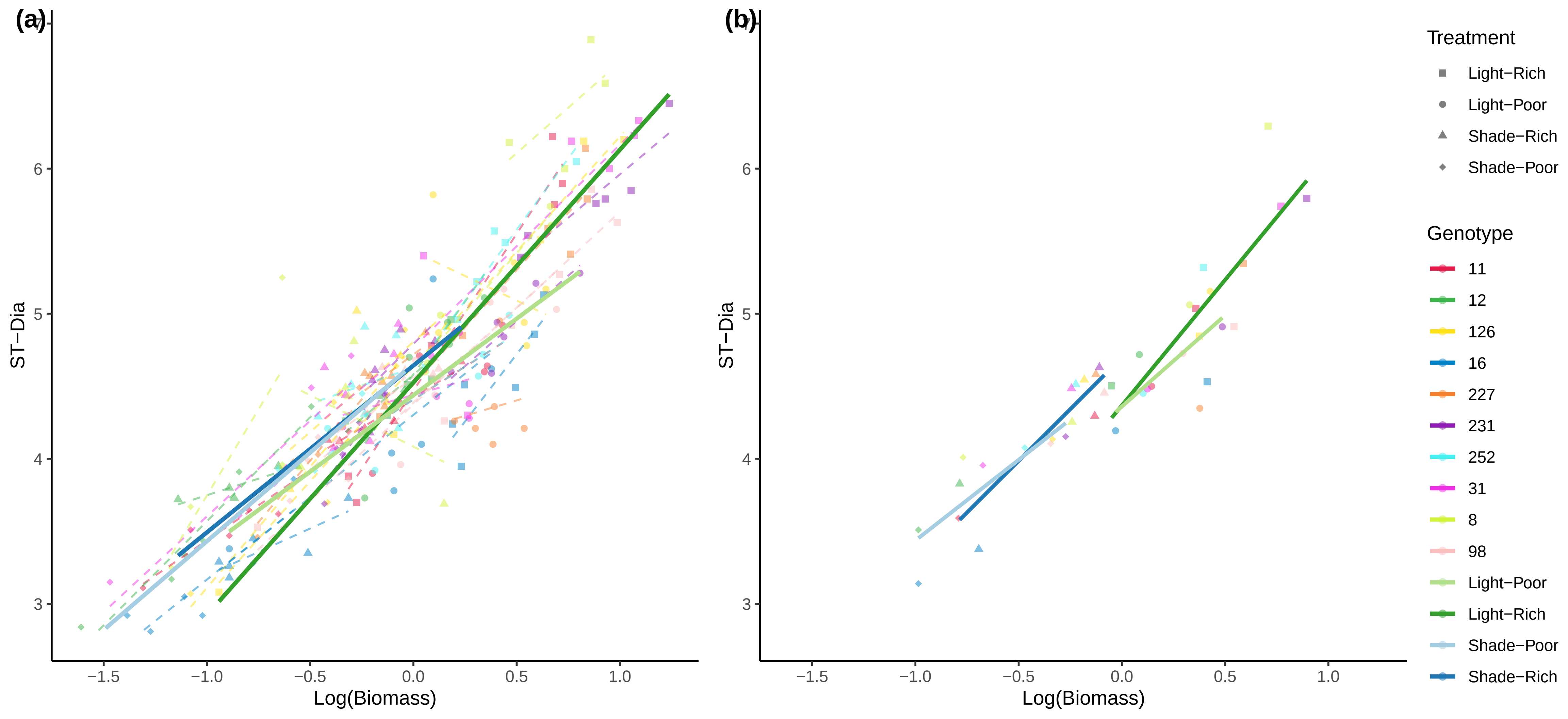

(a)
(b)
ST−Dia
Log(Biomass)
Treatment
Light−Rich
Light−Poor
Shade−Rich
Shade−Poor
Genotype
11
12
126
16
227
231
252
31
8
98
Light−Poor
Light−Rich
Shade−Poor
Shade−Rich

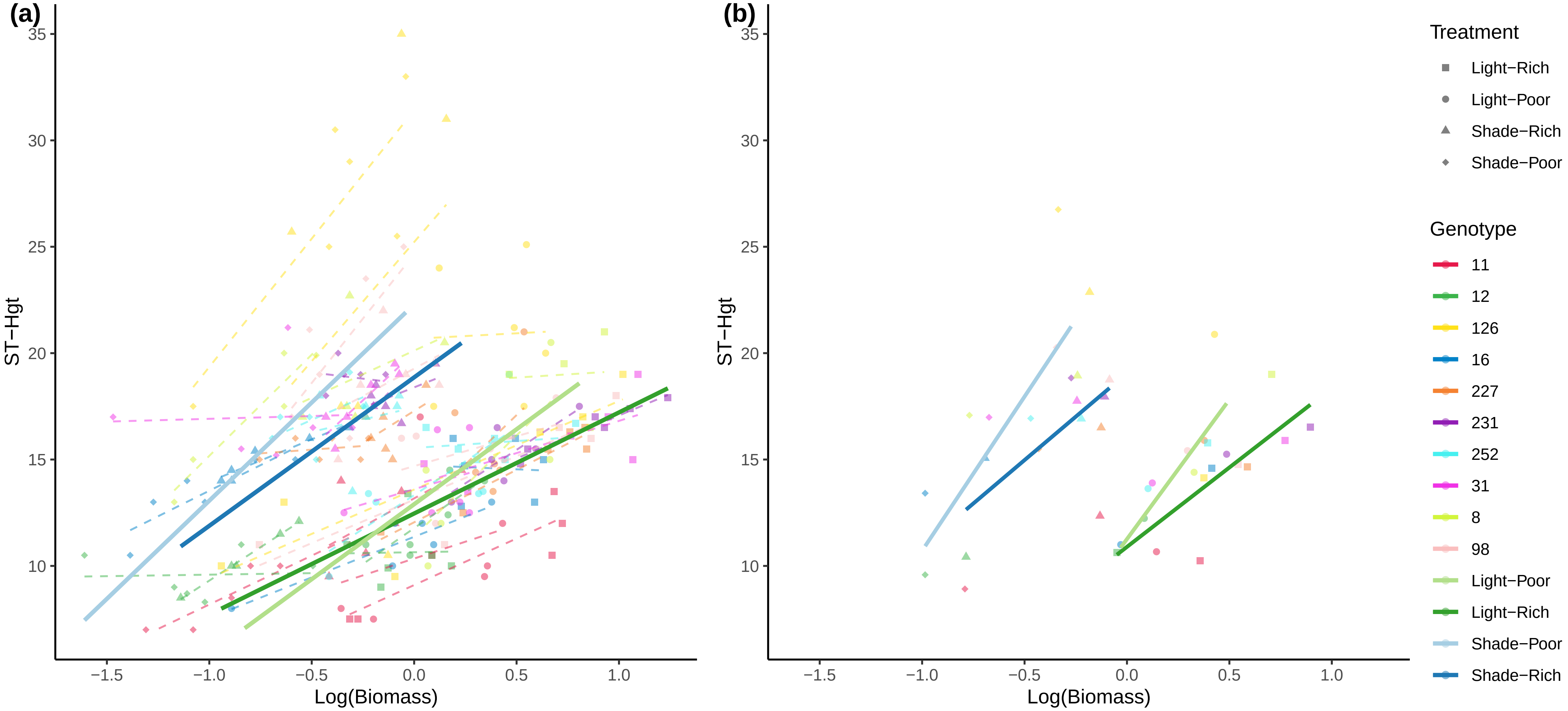

(a)
(b)
ST−Hgt
Log(Biomass)
35
30
25
20
15
10
−1.5
−1.0
−0.5
0.0
0.5
1.0
Treatment
Light−Rich
Light−Poor
Shade−Rich
Shade−Poor
Genotype
11
12
126
16
227
231
252
31
8
98
Light−Poor
Light−Rich
Shade−Poor
Shade−Rich

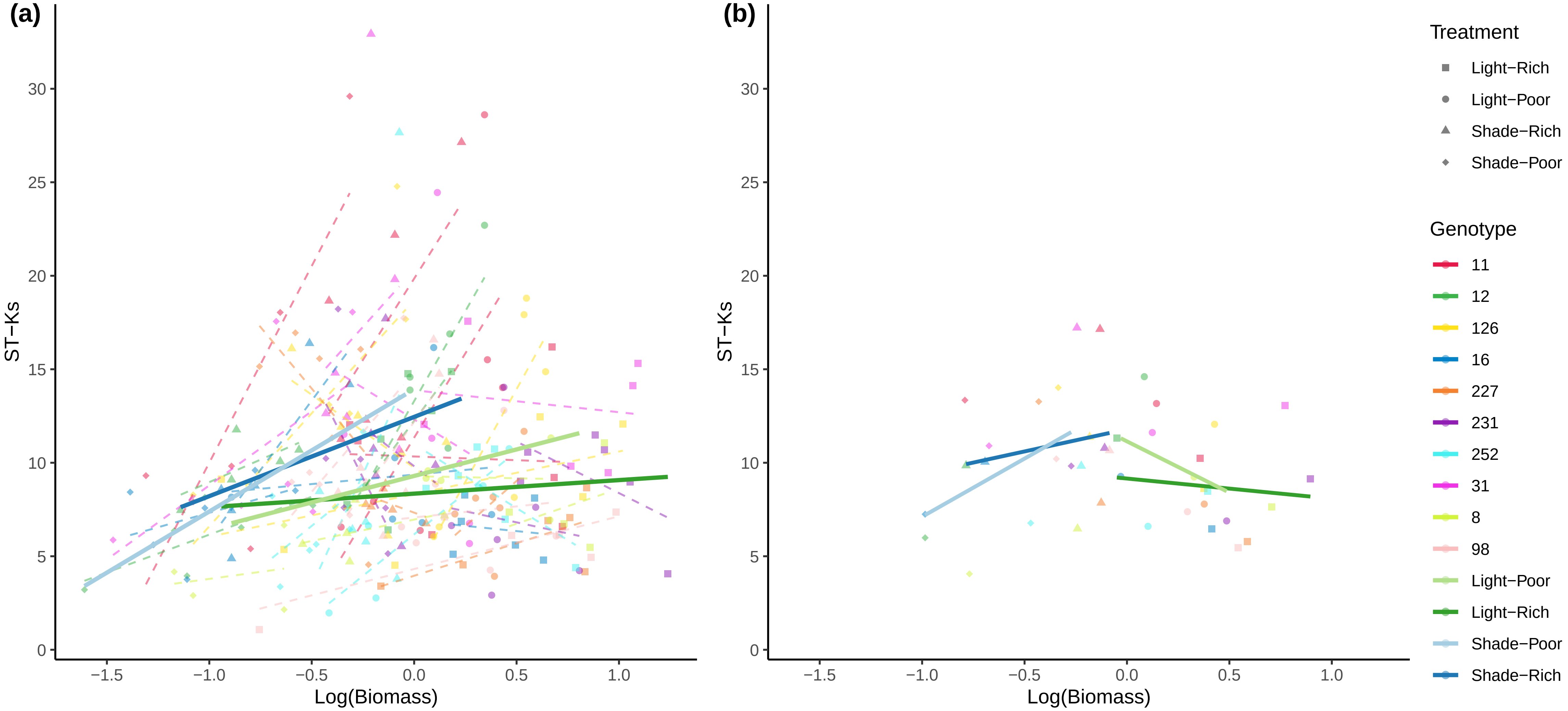

(a)
(b)
ST−Ks
Log(Biomass)
0
5
10
15
20
25
30
−1.5
−1.0
−0.5
0.0
0.5
1.0
Treatment
Light−Rich
Light−Poor
Shade−Rich
Shade−Poor
Genotype
11
12
126
16
227
231
252
31
8
98
Light−Poor
Light−Rich
Shade−Poor
Shade−Rich

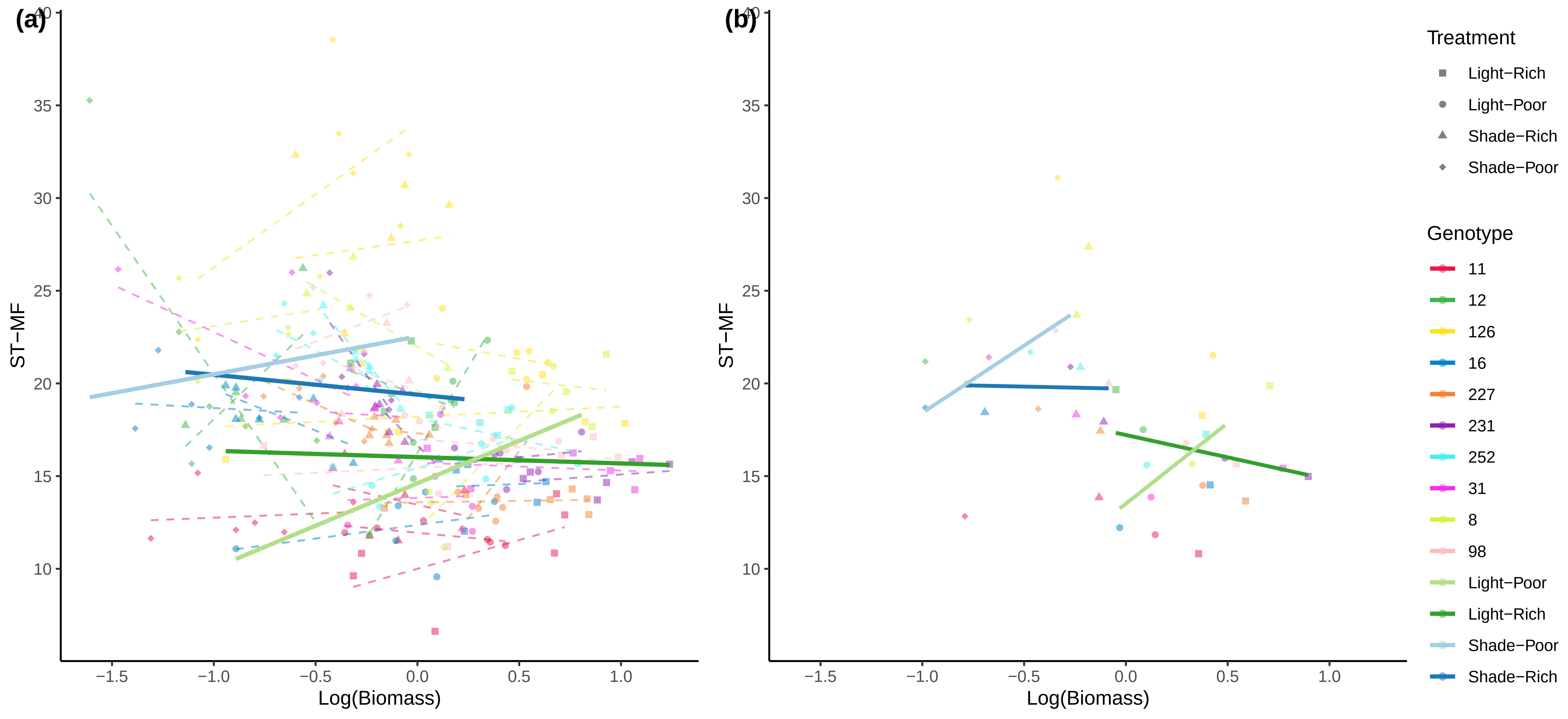
(a)
(b)
ST−MF
Log(Biomass)
Treatment
Light−Rich
Light−Poor
Shade−Rich
Shade−Poor
Genotype
11
12
126
16
227
231
252
31
8
98
Light−Poor
Light−Rich
Shade−Poor
Shade−Rich

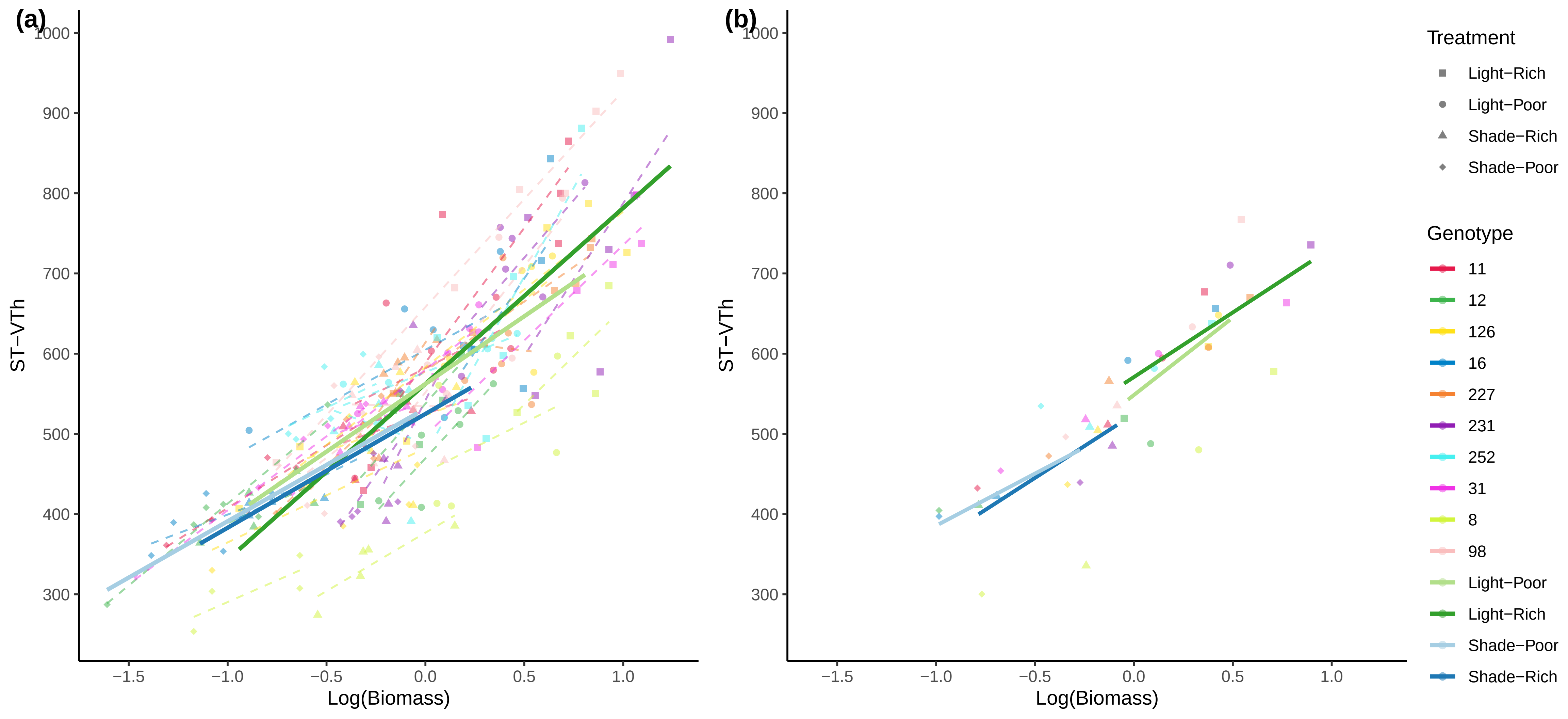

(a)
(b)
ST−VTh
Log(Biomass)
1000
900
800
700
600
500
400
300
−1.5
−1.0
−0.5
0.0
0.5
1.0
Treatment
Light−Rich
Light−Poor
Shade−Rich
Shade−Poor
Genotype
11
12
126
16
227
231
252
31
8
98
Light−Poor
Light−Rich
Shade−Poor
Shade−Rich

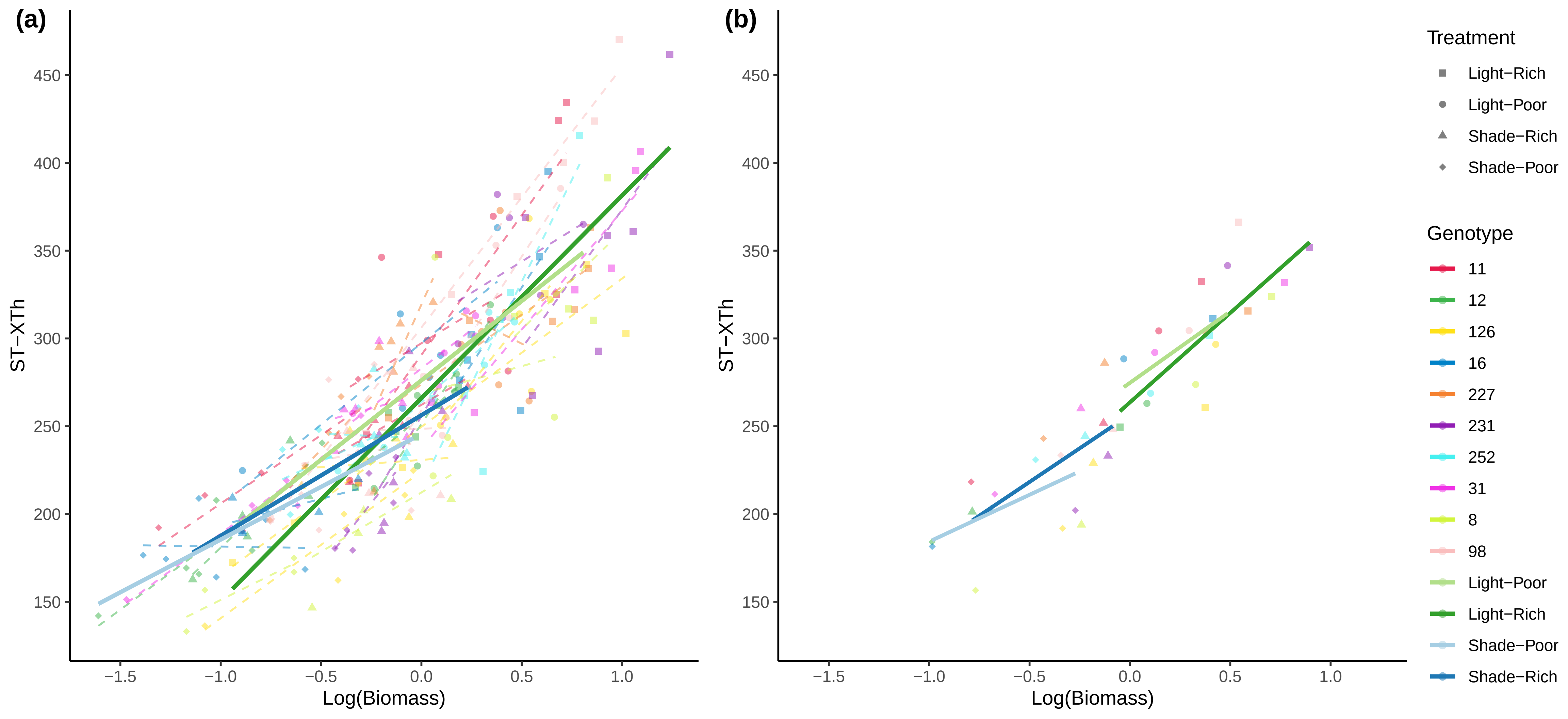
(a)
(b)
ST−XTh
Log(Biomass)
450
400
350
300
250
200
150
−1.5
−1.0
−0.5
0.0
0.5
1.0
Treatment
Light−Rich
Light−Poor
Shade−Rich
Shade−Poor
Genotype
11
12
126
16
227
231
252
31
8
98
Light−Poor
Light−Rich
Shade−Poor
Shade−Rich

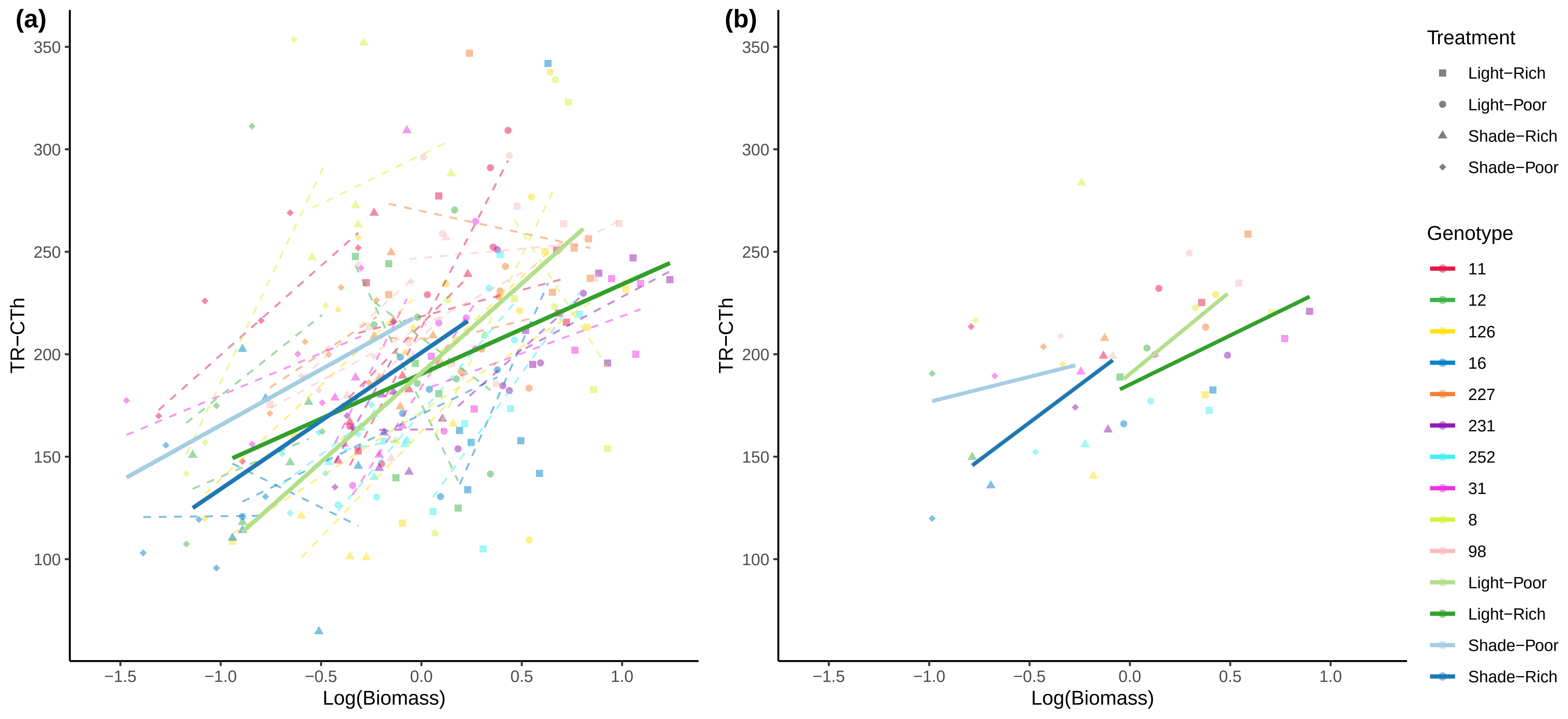
(a)
(b)
350
300
250
200
150
100
TR−CTh
−1.5
−1.0
−0.5
0.0
0.5
1.0
Log(Biomass)
Treatment
Light−Rich
Light−Poor
Shade−Rich
Shade−Poor
Genotype
11
12
126
16
227
231
252
31
8
98
Light−Poor
Light−Rich
Shade−Poor
Shade−Rich

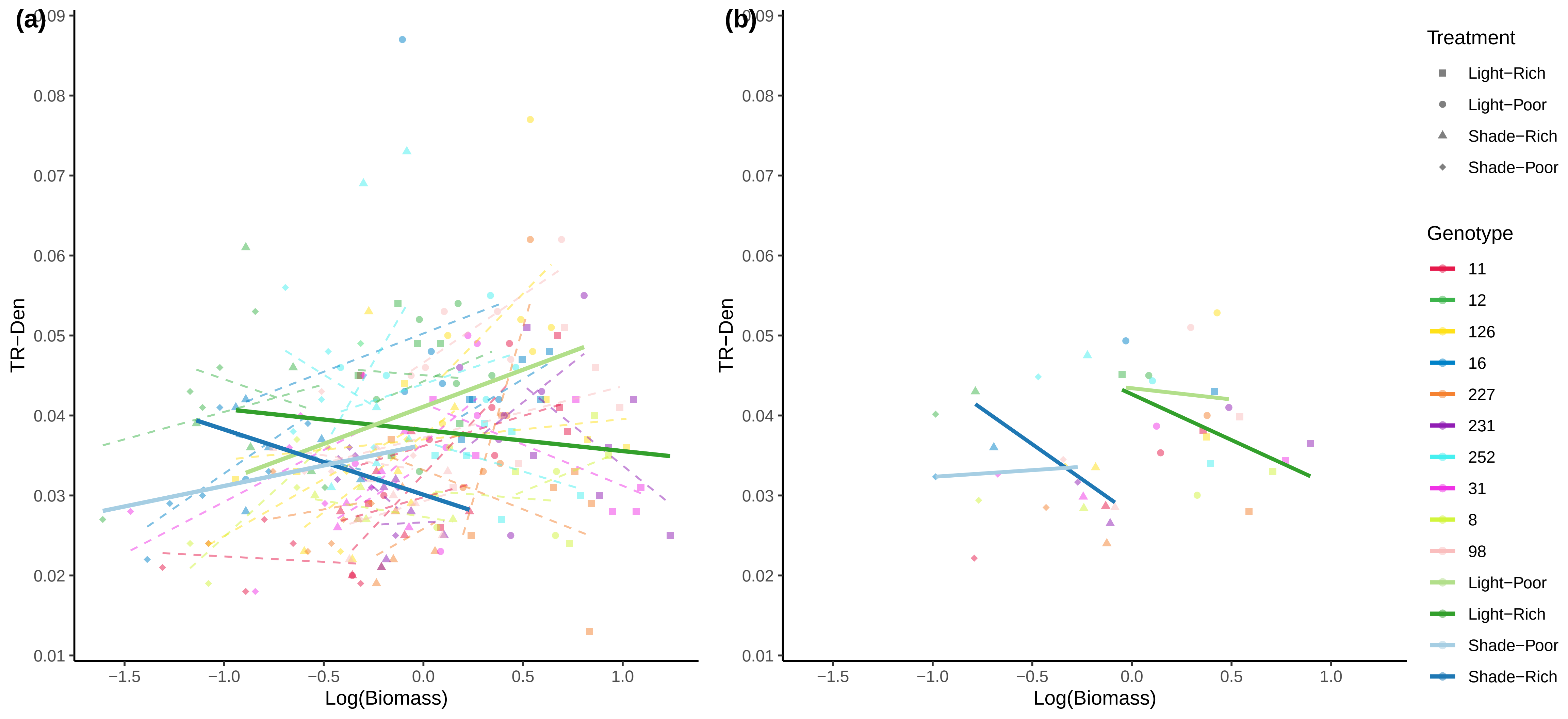

(a)
(b)
0.09
0.08
0.07
0.06
0.05
0.04
0.03
0.02
0.01
TR−Den
−1.5
−1.0
−0.5
0.0
0.5
1.0
Log(Biomass)
Treatment
Light−Rich
Light−Poor
Shade−Rich
Shade−Poor
Genotype
11
12
126
16
227
231
252
31
8
98
Light−Poor
Light−Rich
Shade−Poor
Shade−Rich

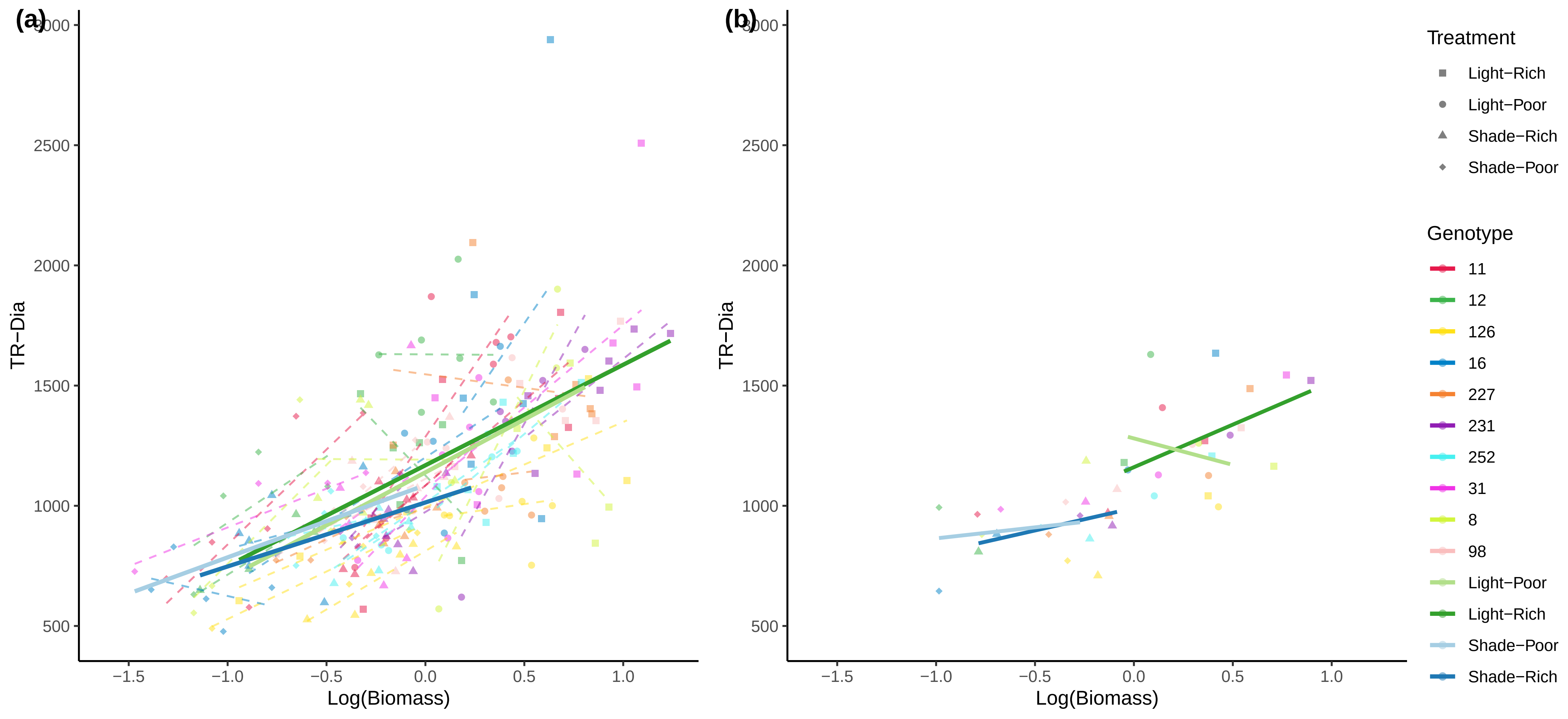

(a)
(b)
3000
2500
2000
1500
1000
500
TR−Dia
−1.5
−1.0
−0.5
0.0
0.5
1.0
Log(Biomass)
Treatment
Light−Rich
Light−Poor
Shade−Rich
Shade−Poor
Genotype
11
12
126
16
227
231
252
31
8
98
Light−Poor
Light−Rich
Shade−Poor
Shade−Rich

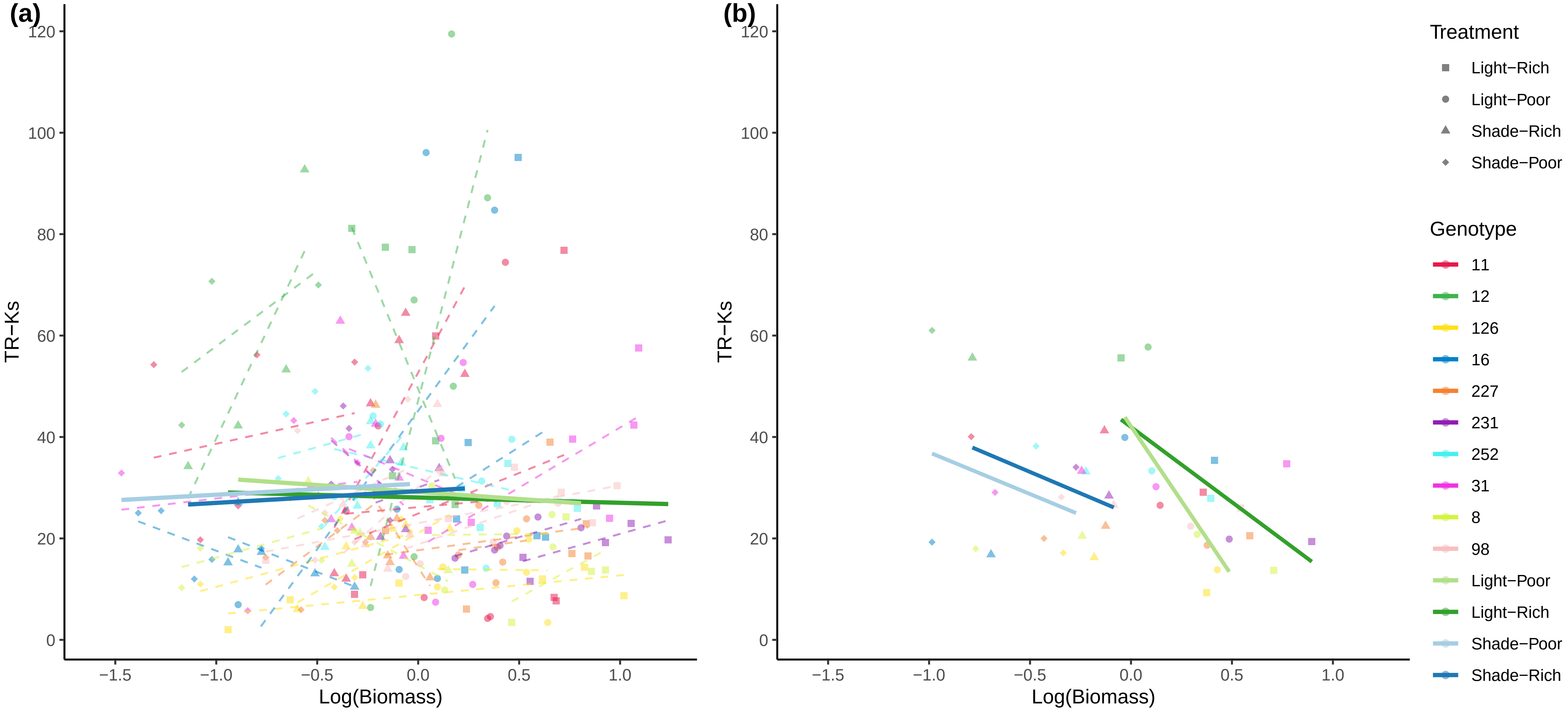

(a)
(b)
TR−Ks
Log(Biomass)
120
100
80
60
40
20
0
−1.5
−1.0
−0.5
0.0
0.5
1.0
Treatment
Light−Rich
Light−Poor
Shade−Rich
Shade−Poor
Genotype
11
12
126
16
227
231
252
31
8
98
Light−Poor
Light−Rich
Shade−Poor
Shade−Rich

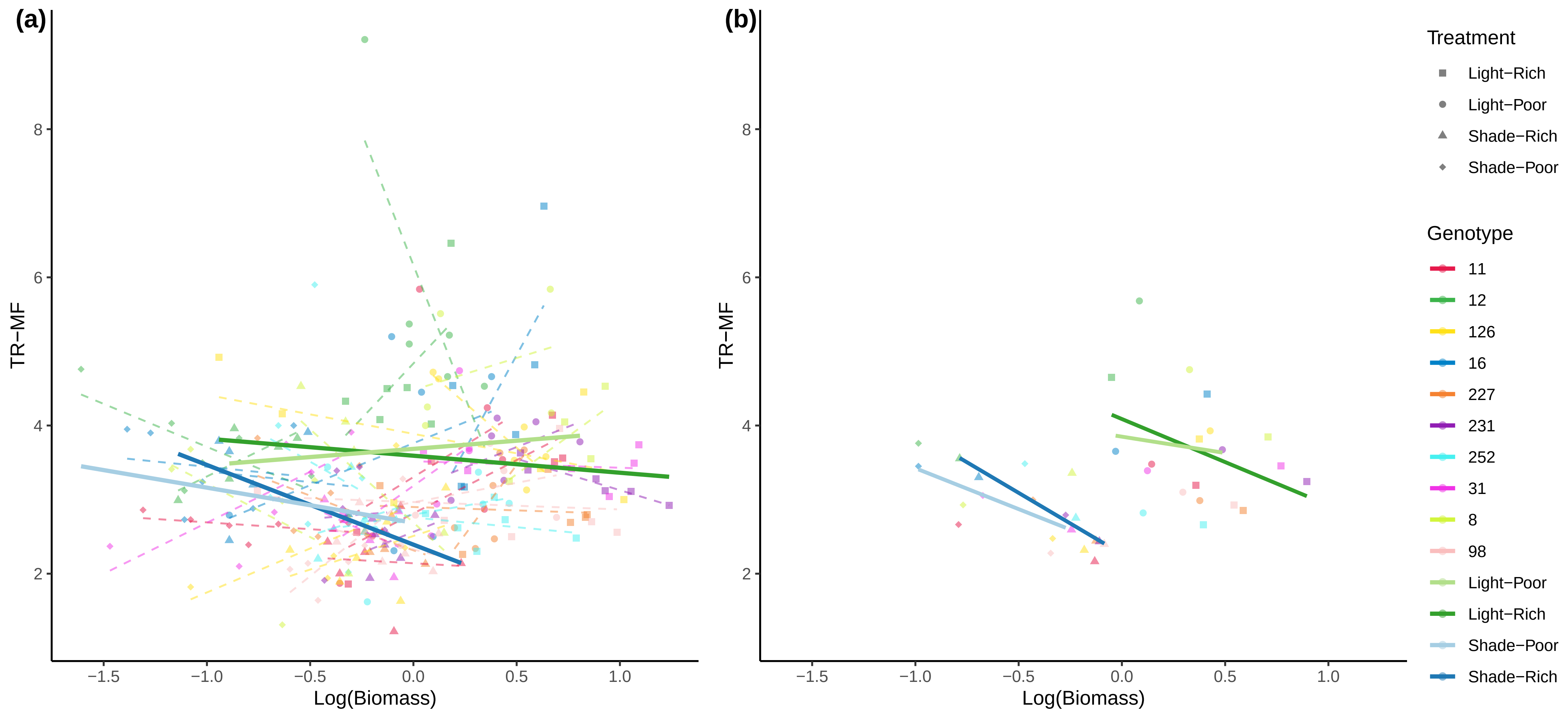
(a)
(b)
TR−MF
Log(Biomass)
Treatment
Light−Rich
Light−Poor
Shade−Rich
Shade−Poor
Genotype
11
12
126
16
227
231
252
31
8
98
Light−Poor
Light−Rich
Shade−Poor
Shade−Rich

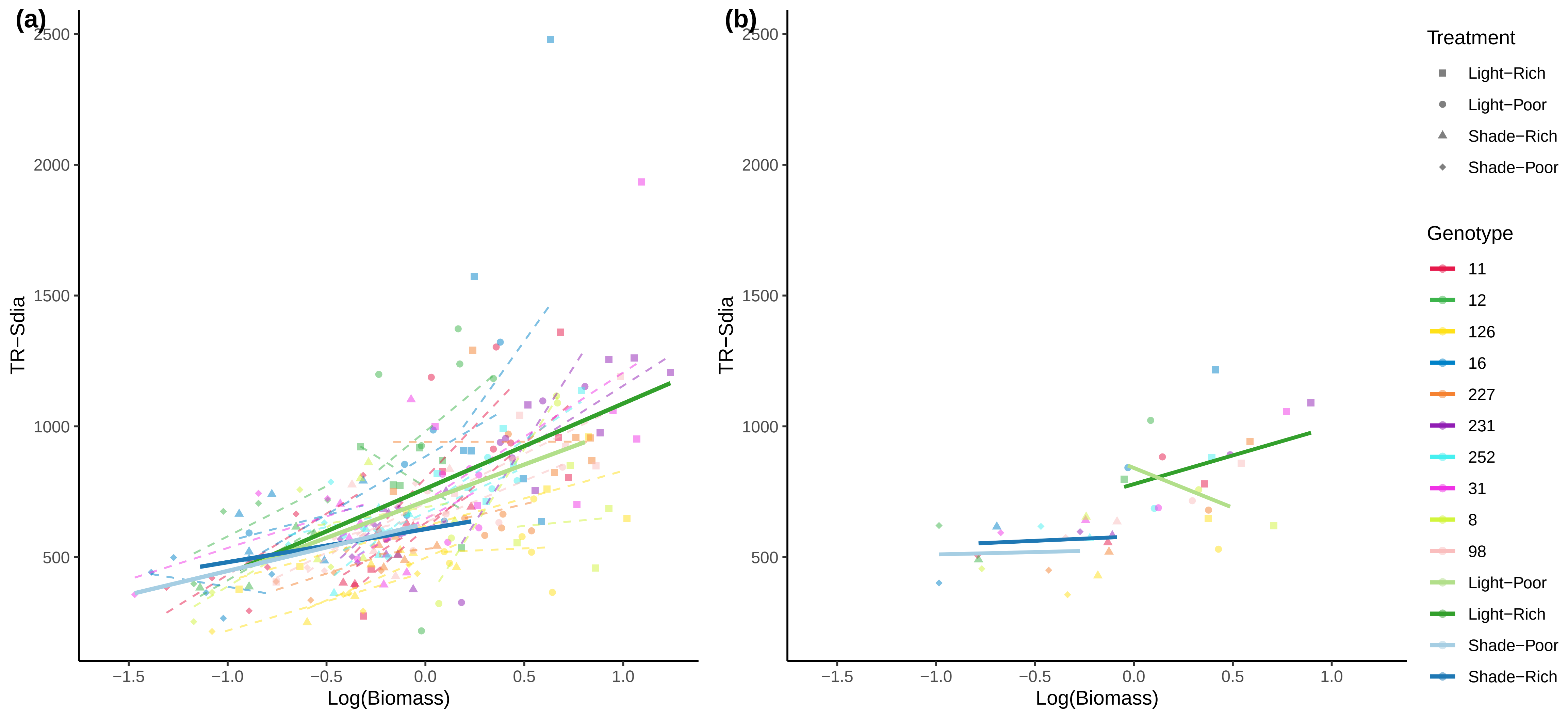
(a)
(b)
TR−Sdia
Log(Biomass)
2500
2000
1500
1000
500
−1.5
−1.0
−0.5
0.0
0.5
1.0
Treatment
Light−Rich
Light−Poor
Shade−Rich
Shade−Poor
Genotype
11
12
126
16
227
231
252
31
8
98
Light−Poor
Light−Rich
Shade−Poor
Shade−Rich

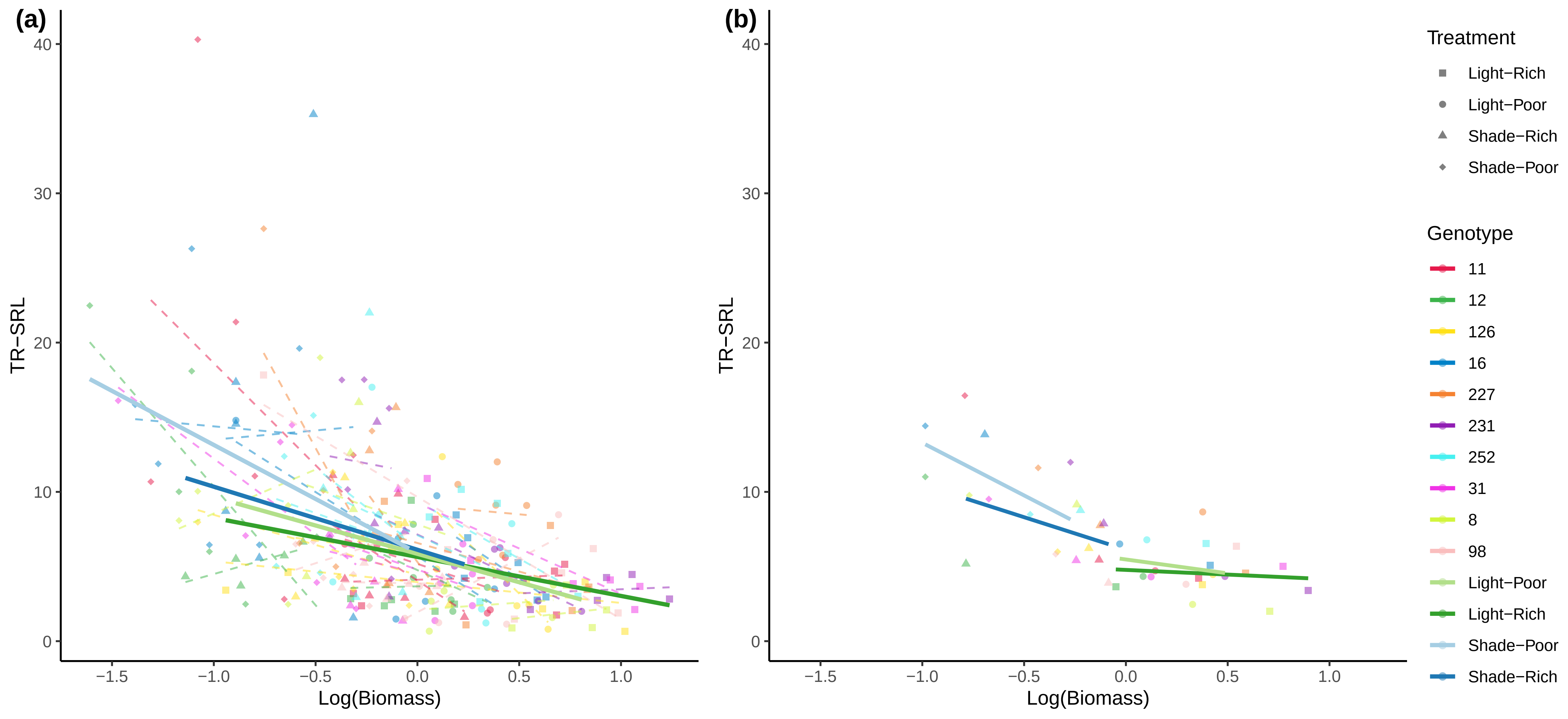

(a)
(b)
TR−SRL
Log(Biomass)
40
30
20
10
0
−1.5
−1.0
−0.5
0.0
0.5
1.0
Treatment
Light−Rich
Light−Poor
Shade−Rich
Shade−Poor
Genotype
11
12
126
16
227
231
252
31
8
98
Light−Poor
Light−Rich
Shade−Poor
Shade−Rich

**Supplemental Figure S4.** Trait contributions to principal components of figure 5.

Trait contributions PC1 Fig 5a

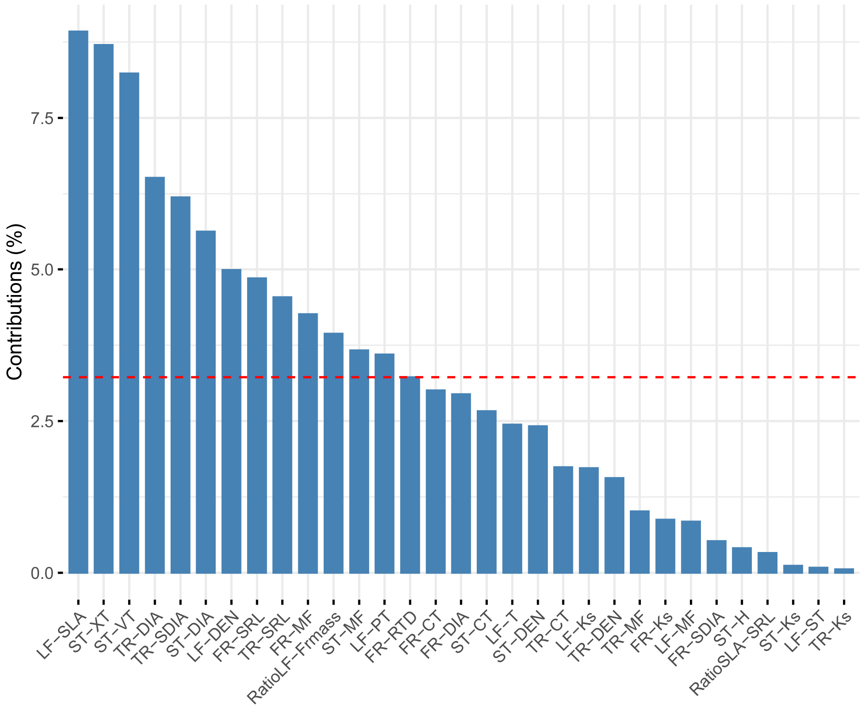


Trait contributions PC2 Fig 5a

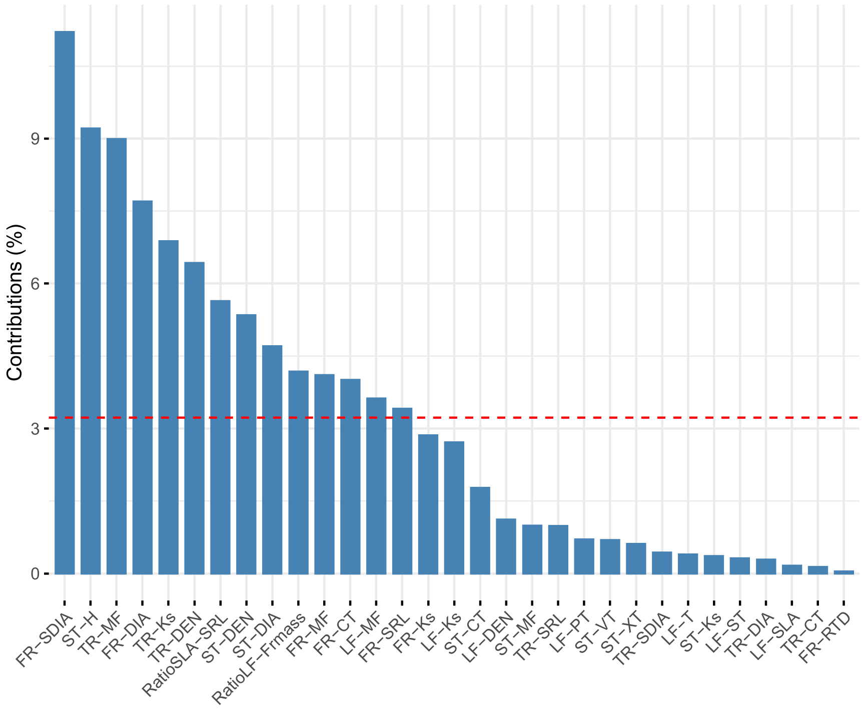

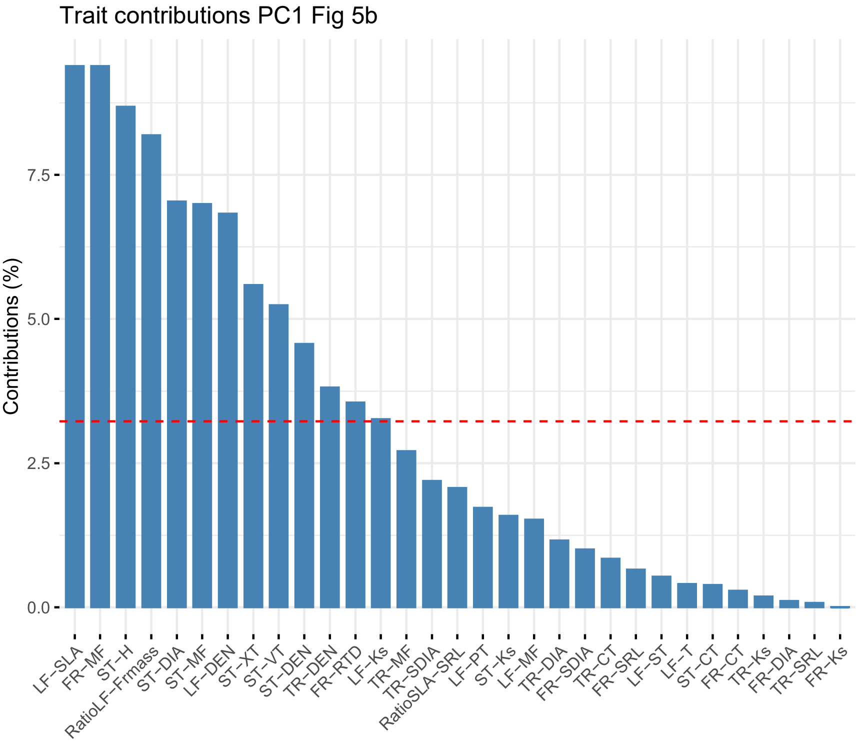
Trait contributions PC1 Fig 5b
Contributions (%)
0.0
2.5
5.0
7.5
LF−SLA
FR−MF
ST−H
RatioLF−Frmass
ST−DIA
ST−MF
LF−DEN
ST−XT
ST−VT
ST−DEN
TR−DEN
FR−RTD
LF−Ks
TR−MF
TR−SDIA
RatioSLA−SRL
LF−PT
ST−Ks
LF−MF
TR−DIA
FR−SDIA
TR−CT
FR−SRL
LF−ST
LF−T
ST−CT
FR−CT
TR−Ks
FR−DIA
TR−SRL
FR−Ks

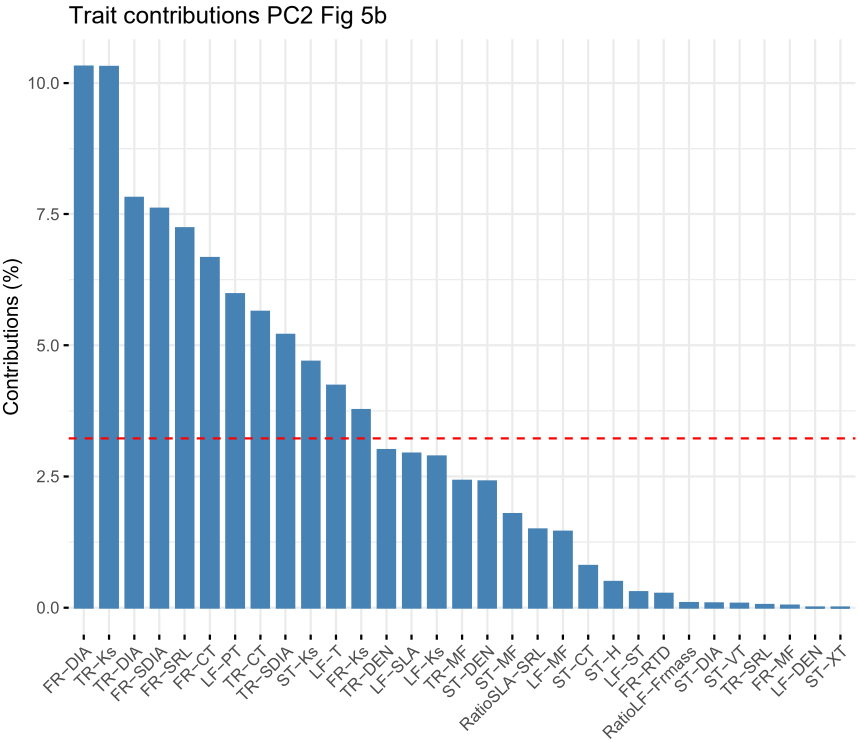
Trait contributions PC2 Fig 5b
Contributions (%)
0.0
2.5
5.0
7.5
10.0
FR−DIA
TR−Ks
TR−DIA
FR−SDIA
FR−SRL
FR−CT
LF−PT
TR−CT
TR−SDIA
ST−Ks
LF−T
FR−Ks
TR−DEN
LF−SLA
LF−Ks
TR−MF
ST−DEN
ST−MF
RatioSLA−SRL
LF−MF
ST−CT
ST−H
LF−ST
FR−RTD
RatioLF−Frmass
ST−DIA
ST−VT
TR−SRL
FR−MF
LF−DEN
ST−XT
